# A crucial role for dynamic expression of components encoding the negative arm of the circadian clock

**DOI:** 10.1101/2023.04.24.538162

**Authors:** Bin Wang, Xiaoying Zhou, Arminja N. Kettenbach, Hugh D. Mitchell, Lye Meng Markillie, Jennifer J. Loros, Jay C. Dunlap

## Abstract

In the *Neurospora* circadian system, the White Collar Complex (WCC) drives expression of the principal circadian negative arm component *frequency* (*frq*). FRQ interacts with FRH (FRQ-interacting helicase) and CK-1 forming a stable complex that represses its own expression by inhibiting WCC. In this study, a genetic screen identified a gene, designated as *brd-8*, that encodes a conserved auxiliary subunit of the NuA4 histone acetylation complex. Loss of *brd-8* reduces H4 acetylation and RNA polymerase (Pol) II occupancy at *frq* and other known circadian genes, and leads to a long circadian period, delayed phase, and defective overt circadian output at some temperatures. In addition to strongly associating with the NuA4 histone acetyltransferase complex, BRD-8 is also found complexed with the transcription elongation regulator BYE-1. Expression of *brd-8, bye-1, histone hH2Az*, and several NuA4 subunits is controlled by the circadian clock, indicating that the molecular clock both regulates the basic chromatin status and is regulated by changes in chromatin.

Taken together, our data identify new auxiliary elements of the fungal NuA4 complex having homology to mammalian components, which along with conventional NuA4 subunits, are required for timely and dynamic *frq* expression and thereby a normal and persistent circadian rhythm.

## INTRODUCTION

Circadian clocks control a wide variety of physiological, biochemical, and behavioral processes in most eukaryotes and certain prokaryotes. At the molecular level, circadian systems are composed of positive and negative arms and the latter repress their own expression through inhibiting the positive arm. In *Neurospora*, the White Collar Complex (WCC) formed by WC-1 and WC-2 transcriptionally activates the pacemaker gene, *frequency* (*frq*), by binding to two promoter elements: *Clock box* (*C-box*) in the dark and *Proximal Light-Response Element* (*pLRE*) in the light^1–3^. FRQ, the *frq* gene product, associates with FRH (FRQ-interacting RNA helicase)^4,5^ and CKI (casein kinase I)^6,7^ to form the FFC complex, promoting WCC phosphorylation at a group of residues^6,8,9^ to repress its activity and thereby close the feedback loop.

*frq* is strongly activated in the light and is rhythmically transcribed and translated in the dark^10^, events that are highly regulated and whose progress impacts timekeeping; *frq* lies in a region of highly accessible chromatin associated with rapid nucleosome turnover when active^11^. In chromatin, nucleosomes impede transcription initiation by Pol II by inhibiting assembly of the preinitiation complex (PIC) at the promoter^12^. The positive charges on lysine residues of histones can be neutralized by acetylation, which weakens histone-DNA interactions and reduces accessibility to other proteins relevant to gene expression^13^. Acetylation also provides recognition sites for chromatin remodeling complexes bearing bromodomains^14^. Nucleosomes can inhibit transcription elongation by Pol II and their displacement at coding sequences directly correlates with the transcription pace of Pol II^15^. The major histone acetyltransferase for histone H4, H2A, and H2A.Z^16–19^ in yeast is the thirteen subunit 1.3 MDa NuA4 (<u>Nu</u>cleosome <u>a</u>cetyltransferase of histone H<u>4</u>) complex, and its catalytic subunit is Esa1. NuA4 is recruited both to gene promoters and coding regions and stimulates the passage of elongating Pol II in transcription initiation and elongation by loosening DNA-histone contacts^20^. Evidence suggests that most histone acetylation at promoter regions precedes and is independent of transcription. In humans, the orthologous Tip60 complex that bears HAT activity is a multiprotein complex with at least 16 subunits^21,22^, and may have arisen as a fusion form of two yeast HAT complexes, NuA4 and SWR, the multi-subunit complex that incorporates histone variant H2A.Z into chromatin^21^. In addition to histones, TIP60 acetylates BMAL1 in the mammalian clock, providing a docking site that brings the BRD4-P-TEFb complex to DNA-bound BMAL1, promoting release of Pol II and elongation of circadian transcripts^23^. The BYpass of Ess1 (BYE1) protein is a

*Saccharomyces cerevisiae* transcription elongation factor^24,25^ whose human homologs are PHF3 and DIDO^26^. BYE-1 interacts with Pol II and can positively or negatively regulate transcription elongation^24–26^. Histone H4 acetylation determines chromatin higher-order structure and functional interactions between remodeling enzymes and the chromatin fibers^27^; disruption of Tip60 or NuA4 and impairment of histone H4 acetylation have been implicated in DNA damage repair, plant developmental processes, behavioral variability, neurodegenerative diseases, cell cycle progression, cancer, and ageing^28–44^.

In this study, we have identified a conserved bromodomain-containing NuA4-associated subunit, BRD-8, which is required for the NuA4 HAT activity on histone H4 at multiple loci including *frq* and is required for normal *frq* expression, period determination, and persistence of rhythmicity. BRD-8 also complexes with an ortholog of Saccharomyces BYE-1, suggesting a connection to transcription elongation. In the absence of *brd-8*, both histone acetylation and Pol II levels at *frq* are sharply reduced, consistent with reduced expression of *frq*. Interestingly, expression of *brd-8*, *bye-1*, and several NuA4 subunits is controlled by the circadian clock, forming a nested feedback loop surrounding the core transcription translation feedback loop. These and prior data are consistent with an important role for robust and timely transcription of the negative arm proteins in the circadian feedback loop and suggest a link between the circadian clock and the basic transcriptional machinery.

## RESULTS

### Loss of *brd-8* impacts the *Neurospora* clock

To identify genes regulating the circadian clock, single gene deletion strains of *Neurospora*^45^ were screened on race tube medium bearing menadione which facilitates visualization of the circadian phenotypes^46^. Two deletion strains lacking the same gene, *ncu09482* (now designated as *brd-8*, see below), displayed an arrhythmic overt conidiation phenotype on race tubes (Figure 1a). To verify the circadian phenotype observed using menadione, Δ*brd-8* was backcrossed to the *ras-1^bd^* and *frq C-box*-driven *luciferase* reporters that have been widely used to visualize overt rhythms and continuously report changes in WCC activity within the core clock respectively. Δ*brd-8* in the *ras-1^bd^*background showed a ∼3 hour longer period with a slightly reduced growth rate for the first three days and subsequently became arrhythmic by both race tube and luciferase assays (Figure 1a and Supplementary Figure 1). Consistent with this, the robustness of the *frq C-box*-driven *luciferase* reporter was also dramatically impaired after three days in Δ*brd-8* (Figure 1a and Supplementary Figure 2). The long period phenotype was not rescued when medium having a lower concentration (0.01%) of glucose (Supplementary Figures 1 and 2) was used to reduce conidiation and thereby better visualize the overt rhythm; a ∼3-hr phase delay was also observed in Δ*brd-8* (Figure 1a), and the reduced *frq C-box*-driven bioluminescence signal is consistent with reduced *C-box*-driven expression which would lead to reduced *frq* expression. Circadian clocks maintain their period lengths across a range of physiological temperatures. To test the performance of Δ*brd-8* at different temperatures, circadian periods of WT and Δ*brd-8* were examined at 20°, 25°, and 30 °C (Figure 1b); Δ*brd-8* displays arrhythmic overt phenotypes at 30 °C in media bearing either a high (0.1%) or low (0.01%) percentage of glucose (Supplementary Figure 1), whereas it is rhythmic at all temperatures tested by luciferase analyses (Supplementary Figure 2), suggesting that *brd-8* also controls the circadian output at high temperature independent of its role in regulating the period length.

**Figure 1.**
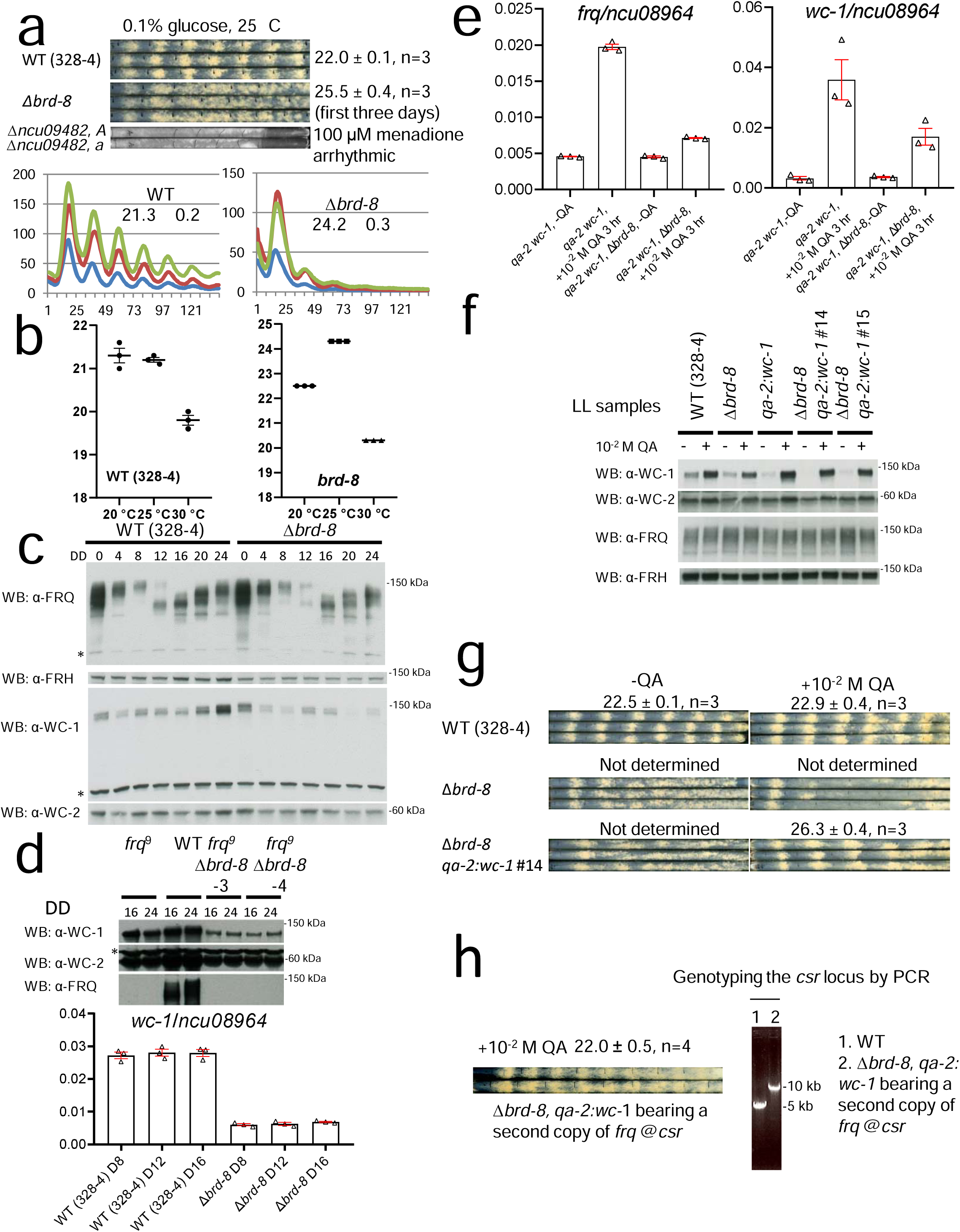
Identification of *brd-8* as a regulator of the *Neurospora* core circadian oscillator and output. Δ*brd-8* was assayed by race tube (**a**, upper) and luciferase (**a**, lower) analyses. Of the race tube analyses, only Δ*ncu09482*, *a* and Δ*ncu09482*, *A* were grown on regular race tube medium containing 100 μM menadione and showed an arrhythmic overt clock. (**b**) Circadian periods of WT and Δ*brd-8* at 20°, 25°, and 30 °C were determined by the luciferase assay. The period data presented here are the average +/-the standard error of the mean (SEM). Raw data are shown in Supplementary Figure 2. (**c**) Expression of FRQ, FRH, WC-1, and WC-2 in WT and Δ*brd-8* over 24 hr. DD, hours after the light to dark transfer. The experiment was repeated three times with similar results. (**d**) Upper panel, total WC-1, WC-2, and FRQ were followed by Western blotting at DD16 and DD24 in WT, *frq^9^*, and *frq^9^*; Δ*brd-8*. Lower panel, mRNAs extracted from samples cultured in the dark for 8, 12, and 16 hrs. (**e**) Expression of *wc-1* and *frq* in *qa-2*:*wc-1* and *qa-2:wc-1*; Δ*brd-8* was determined by RT-qPCR in the absence or presence of 10^-2^ M QA for 3 hrs. In Figures 1d and 1e, the RT-qPCR data are representatives of three biological experiments (n=3) and reported as the average +/-SEM; gene expression was normalized to that of *ncu08964*. (**f**) Total WC-1, WC-2, FRQ, and FRH were measured in WT, Δ*brd-8*, *qa-2:wc-1*, and *qa-2 wc-1*; Δ*brd-8* with or without QA in the medium as indicated. The experiment was done three times, and similar results were obtained. (**g**) Race tube analyses of WT, Δ*brd-8*, *qa-2:wc-1*, and *qa-2*:*wc-1* in the absence or presence of 10^-2^ M QA (**h**) Knocking in a second copy of *frq* at the *csr* locus rescues the long period length observed in Δ*brd-8*. The amplicon with a primer set used in genotyping is against the *csr* locus; in the *frq* knock-in strain, a PCR product with a larger size was amplified by PCR compared to that in the WT. Source data are provided as a Source Data file.

**Figure 2.**
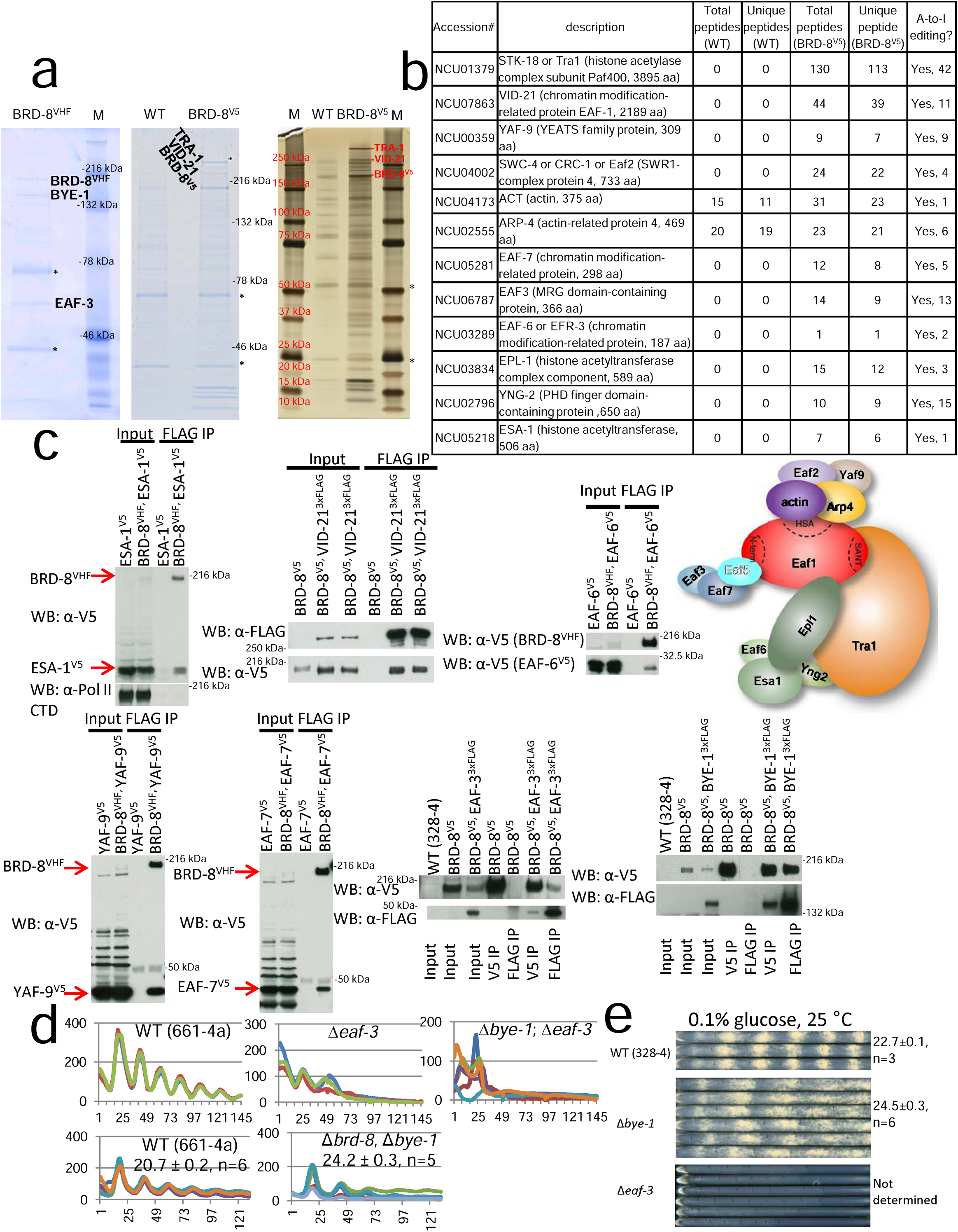
BRD-8 interacts with NuA4 subunits and BYE-1. (**a**) Coomassie blue-and silver-stained gels showing BRD-8^VHF^ or BRD-8^V5^ and its interactome purified from a culture grown in the light. BRD-8 and interactors were affinity-purified, run on SDS-PAGE gels, and analyzed by mass spectrometry. For the BRD-8^VHF^ interactors (middle gel, purified from 30 g of tissue grown in the light by tandem V5, His, and FLAG steps), individual interactor bands were removed for mass spectrometry analyses while for BRD-8^V5^ (left and right gels, each purified from 5 g of tissue grown in the light by a single V5 step), either individual bands or the whole interactome (the entire Coomassie blue-stained gel lane; see Methods) was removed and then analyzed. The rightmost gel was loaded with extracts equivalent to the middle gel but was silver-stained to afford greater sensitivity in visualizing protein bands. Purification with BRD-8^VHF^ was done twice and similar results were seen; interactor identification from BRD-8^V5^ was performed three times, and similar band profiles were observed. (**b**) List of NuA4 subunits interacting with BRD-8^V5^ identified by mass spectrometry (See Supplementary Data 1 for the full list of BRD-8^V5^ interactors identified). The number of A-to-I editing events on these NuA4 subunits (retrieved from ^51^) are also shown in the table. Annotations for the genes were obtained from FungiDB (https://fungidb.org/fungidb/app) and Uniprot (https://www.uniprot.org/) (**c**) Verification of interaction between BRD-8 and NuA4 subunits by co-immunoprecipitation. BRD-8 and individual interactors were tagged and immunoprecipitated as indicated. All epitope tags were added to genes at their native loci except for *eaf-3^3^ ^x^ ^FLAG^* that was targeted at the *csr* locus. The schematic architecture of *Saccharomyces cerevisiae* NuA4 was modified from^95^ with subunit EAF5 dimmed to indicate that it was not found in our extracts. These experiments were carried out three times, and results were similar. (**d**) Bioluminescent analyses of *frq C-box* activity in WT, Δ*eaf-3*, and Δ*eaf-3*; Δ*bye-1* at 25 °C by the luciferase assay. (**e**) Race tube analyses of WT, Δ*eaf-3*, and Δ*bye-1*. Source data can be found in the Source Data file.

In addition to its role as the major transcription factor for *frq*, WC-1 is also the principal blue-light photoreceptor for the organism, directly or indirectly mediating light-induced gene expression^47^. Interestingly, the Δ*brd-8* strain that displayed a ∼25-hr period and delayed dark FRQ expression retains its capacity to detect light to induce expression of light-responsive genes (Supplementary Figures 3). Unexpectedly, light-induced expression of some genes (e.g. *frq*, *vvd*, *csp-1*, and *sub-1*) is even significantly higher in Δ*brd-8* than in WT, consistent with differing roles for BRD-8 depending on the promoter element (*C-box* vs. *pLRE*) used.

### FRQ and WC-1 levels are reduced in the mutant in the dark

To understand the role of BRD-8 at a mechanistic level, core circadian component expression was examined by Western blot (Figure 1c). FRH and WC-2 levels are normal or slightly reduced in Δ*brd-8,* and consistent with the long period of Δ*brd-8*, new FRQ in Δ*brd-8* appears later than that in WT and is slightly reduced. However, the level of WC-1 in Δ*brd-8* is significantly reduced. Active WC-1 is unstable and FRQ, by inactivating WCC, stabilizes WC-1^5,6^, so to test whether the low WC-1 level is caused by a defective negative arm, Δ*brd-8* was backcrossed to the loss of function allele *frq^9^*; WC-1 levels are further reduced in the *frq^9^*; Δ*brd-8* double mutant (Figure 1d), which indicates that *brd-8* regulates WC-1 expression at least partially independently of the negative feedback loop. Reduced WC-1 levels as seen in Δ*brd-8* can result in modestly long circadian periods^48^. To test whether the long circadian period in Δ*brd-8* is caused by the reduced WC-1 level, the native promoter of *wc-1* in

Δ*brd-8* was replaced by the strongly inducible *qa-2* promoter. In the presence of inducer (quinic acid, QA), the *wc-1* and WC-1 levels are raised to that in WT (Figures 1e and 1f) or even higher, but the period defect of Δ*brd-8* is not rescued (Figure 1g), indicating that the longer period in the mutant is not caused by the low WC-1 level. Similarly, to test whether changes in FRQ levels underlie the long period of Δ*brd-8,* a second copy of the *frq* gene with its native promoter was inserted at the *csr* locus of Δ*brd-8*. The long period of Δ*brd-8* was rescued to WT (Figure 1h), indicating that reduced *frq* expression in Δ*brd-8* is the cause of the long period length.

### BRD-8 is a nonessential subunit of the histone acetyltransferase NuA4 complex and interacts with elongation factor BYE-1

To understand how loss of *brd-8* impacts the circadian clock, BRD-8 was C-terminally epitope-tagged with V5, 10 x histidine, and 3 x FLAG (VHF tag) and purified from protein extracts; three distinct protein bands from the stained gel removed and analyzed by tandem mass spectrometry proved to be proteins encoded by NCU09482 (BRD-8, see below), NCU06787 (ortholog of the NuA4 subunit EAF-3), and NCU05943 (ortholog of the regulator of transcription elongation BYE-1^24^), (left gel, Figure 2a). To recover more interactors and particularly those with enzymatic activities, NCU09482 (BRD-8) tagged with V5 was isolated through a single anti-V5 immunoprecipitation step so as to preserve weak interactions (middle and right gels, Figure 2a), and the proteins analyzed by tandem mass spectrometry (MS/MS). Compared with the negative (untagged) control, the NCU09482 (BRD-8)-interactome is highly enriched for subunits of the NuA4 histone acetyltransferase complex (Figure 2b) (Supplementary Data 1). Unlike other subunits of NuA4 that have no peptides recovered from the negative control (WT [untagged] in Figure 2b), Arp4 itself seems a sticky protein with respect to the affinity resin, similar to actin in the same figure. In *Neurospora*, Arp4 may not exist only in the NuA4 complex but may also be present as an abundant noncomplexed protein that can bind to the antibody-conjugated resin in a nonspecific manner. To verify these interactions *in vivo*, NCU09482 (BRD-8) was C-terminally tagged with V5 or VHF while BYE-1 and most NuA4 subunits including VID-21, EAF-6, EAF-3, YAF-9, EAF-7, ESA-1 etc. (Figure 2c), were individually tagged with 3 x FLAG or V5. Immunoprecipitation using V5 or FLAG antibody confirmed the interaction between BRD-8 and all NuA4 subunits as well as BYE-1 *in vivo* (Figure 2c). NGF-1, a histone H3 acetyltransferase and the ortholog of Saccharomyces GCN5, was chosen as a control to prove the specificity of interactions between BRD-8 and its interactors. No interaction was detected between BRD-8 and NGF-1 (Supplementary Figure 4a), indicating the interaction between BRD-8 and NuA4 subunits is not through promiscuous binding with whole chromatin; from the interactor list identified with BRD-8^V5^ (Supplementary Data 1), fatty acid synthase beta subunit (CEL-2) and HSP-70-5 (HSP70-like) were selected as controls to validate the mass spectrometry result and their interactions with BRD-8 were also not seen (Supplementary Figures 4b and 4c). Because *Neurospora* is an ascomycete fungus, BLASTP was used to search for orthologs to the protein encoded by NCU09482 among fungi; surprisingly no orthologs were detected in Saccharomyces or *S. pombe* although outside of these groups and within the Pezizomycotina, the protein encoded by NCU09482 was found to be well conserved (Supplementary Figure 5). However, NuA4 subunits have not been previously described among these organisms, so the search for orthologs was extended to animals. Interestingly, the top BLASTP hit in humans is BRD8, a component of the animal NuA4 complex not previously found in fungi^49^, and reciprocally BLASTP of the *Neurospora* proteome using human BRD8 identifies the protein encoded by NCU09482 as the next to top hit. Because the protein encoded by NCU09482 specifically associates with NuA4 complex subunits, has a bromodomain, and shows reciprocal (close to) best homology with BRD8, in keeping with prior nomenclature precedents^(e.g. [50])^, it is designated as BRD-8 (bromodomain protein-8).

**Figure 3.**
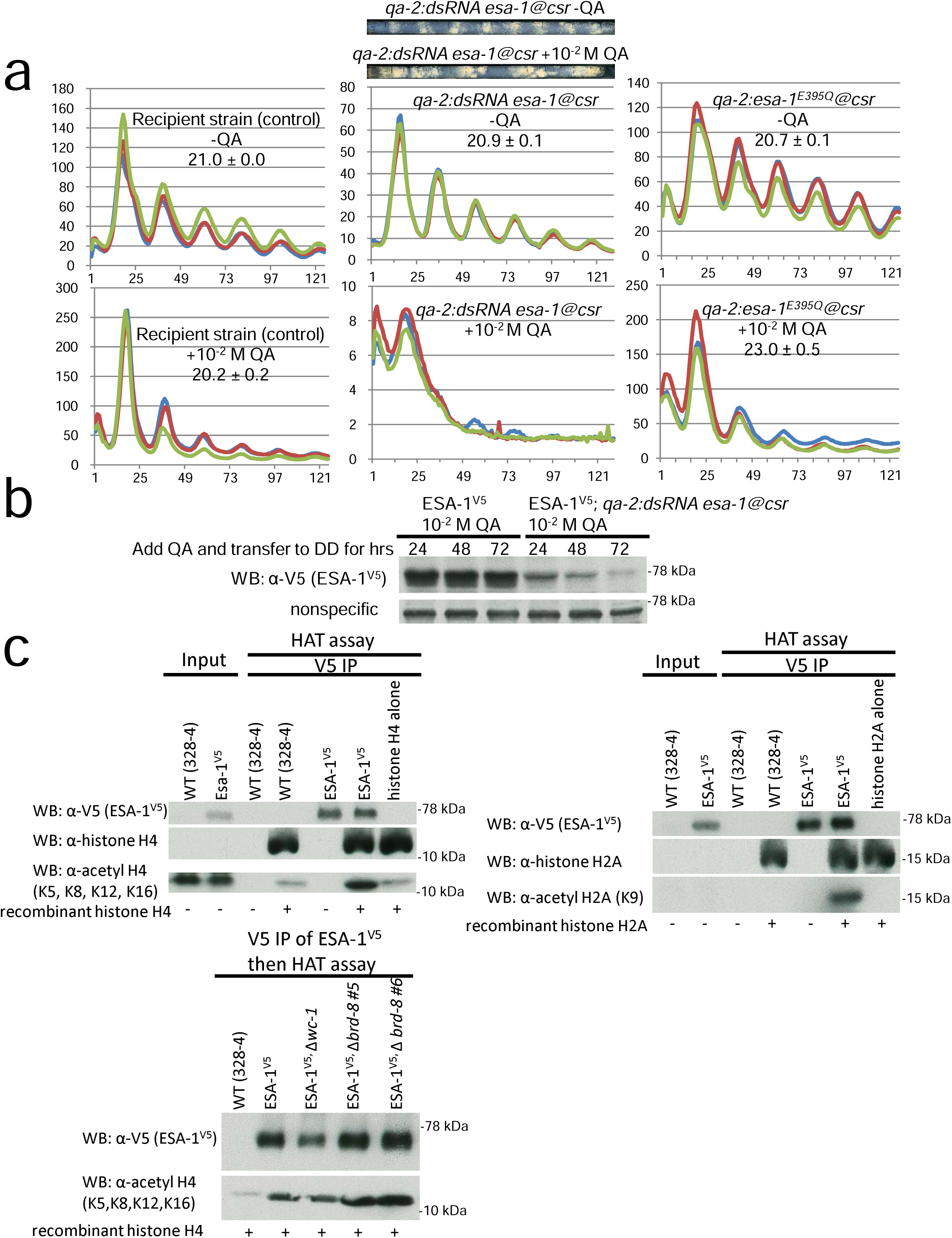
*esa-1* knockdown causes a long period. (**a**) Bioluminescent analyses of strains that knock down *esa-1* (middle) or overexpress *esa-1^E395Q^* (right). *qa-2-*driven double-strand RNA against *esa-1,* or *esa-1^E395Q^* targeting the *csr* locus, was transformed to a WT strain. Circadian periods of WT and Δ*brd-8* at 25 °C were determined by the luciferase assay in the absence or presence of 10^-2^ M QA as indicated. Race tube images on the top indicate the viability of the *qa-2-*driven double-strand RNA against *esa-1* strain grown with 10^-2^ M QA despite a slow and unstable growth rate compared with the no QA control. (**b**) *qa-2:dsRNA* against *esa-1* was transformed to *esa-1^V5^*; strains were cultured in constant light, and 10^-2^ M QA was added to the cultures and immediately transferred to the dark for 24, 47, and 72 hrs; Western blotting against V5 was performed and a non-specific band was shown for equal loadings. The assay was performed three times, and similar results were observed. (**c**) *In vitro* HAT assays of recombinant histone H4 (upper left) or H2A (upper right) with affinity-purified ESA-1^V5^. Details for this assay are described in Methods. Bottom, a HAT assay of ESA-1^V5^ *in vitro*. ESA-1^V5^ in the background of WT, Δ*wc-1*, or Δ*brd-8* was immunoprecipitated by V5 antibody and assayed *in vitro* with recombinant histone H4 to test its histone acetylation activity. Western blotting using antibodies against V5, acetyl H2A (K9), and acetyl H4 (K5, K8, K12, and K16) respectively were followed to show ESA-1^V5^ and acetylated histone H2A and H4 levels. The assay was done three times with similar results. Source data were deposited in the Source Data file.

**Figure 4.**
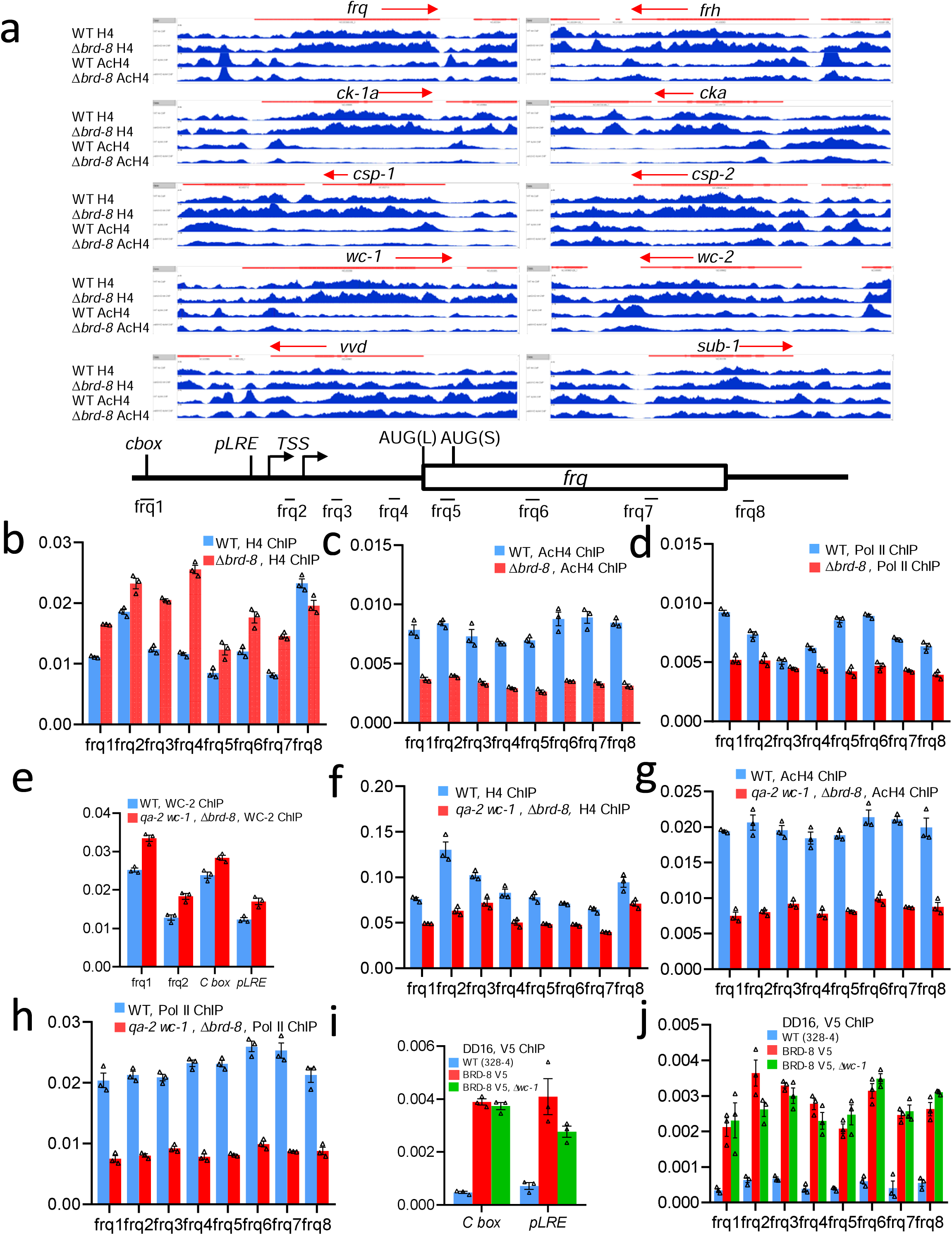
Δ*brd-8* has reduced acetyl-histone H4 and Pol II levels at *frq*. (**a**) The Integrative Genomics Viewer (IGV) visualization of ChIP sequencing data. Antibodies against histone H4 or acetyl histone H4 (AcH4) were applied in ChIP experiments using cultures collected at DD20 for WT and DD26 for Δ*brd-8* to compensate the 3-hr period lengthening and 3-hr phase delay seen in the mutant. Only the ChIP sequencing data are shown. Red bars represent coding regions of the genes: ORFs are in bold while UTRs or introns are in thin bars and arrows designate the direction of transcription. Bottom, diagram of the *frq* gene showing regions amplified by primer sets used in ChIP-PCRs (Figures 4 b-J) as derived from a previous publication^92^ with removal of unrelated information. To validate the data in Figure 4a, independent *Neurospora* samples were collected in the same way as in Figure 4a (WT at DD20 while Δ*brd-8* at DD26), and then ChIP-quantitative PCRs were carried out using histone H4 (**b**) acetyl-histone H4 (**c**), or Pol II (**d**) antibody and *frq*-specific primer sets in (**a**) with WT and Δ*brd-8*. In another set of biologically independent experiments (e-h) to rescue the decreased level of WC-1 in Δ*brd-8*, the promoter of *wc-1* in Δ*brd-8* was replaced by an inducible *qa-2* promoter. Two strains, WT and *qa-2* promoter-driven *wc-1* in Δ*brd-8*, were cultured in the presence of 10^-2^ M QA throughout and afterwards both harvested at DD16. (**e**) ChIP-PCRs were carried out using WC-2 antibody with primers “*frq1*”, “*frq2*”, “*C-box*”, and “*pLRE*” in WT and *qa-2:wc-1*; Δ*brd-8* at DD16. ChIP-PCRs were performed with histone H4 (**f**), acetyl-histone H4 (**g**), or Pol II (**h**) antibody and *frq*-specific primer sets in (**a**) with WT and *qa-2:wc-1*; Δ*brd-8* at DD16. Quantitative ChIP-PCRs were done using V5 antibody with primers “*C-box*” and “*pLRE*” (**i**) and “*frq1-8*” (**j**) in WT, *brd-8^V5^*, and *brd-8^V5^*/Δ*wc-1* at DD16. For ChIP-qPCR assays, three technical replicates were performed, and the bars represent average values plotted as a percentage of the input, with error bars representing the SEM (n= 3). Source data were supplied in the Source Data file.

If loss of BRD-8 impacts NuA4 function leading to period lengthening, then loss of other NuA4 subunits or associated proteins might be expected to have similar and additive effects. This was tested; Δ*eaf-3* shows an impaired long period core clock by a luciferase assay (Figure 2d) and a strong growth defect by race tube analysis (Figure 2e). Δ*bye-1* similarly displays a circadian period lengthened by ∼2 hrs (Figure 2e), while disruption of both *eaf-3* and *bye-1* leads to low and arrhythmic luciferase signals (Figure 2d). The Δ*brd-8,* Δ*bye-1* strain bearing deletions in both components shows a period lengthened to the same degree as the single mutants consistent with them working through the same pathway (Figure 2d). Interestingly, in the perithecia, transcripts of all NuA4 subunits as well as BYE-1 are subject to multiple A-to-I editing events (Figure 2b)^51^. This is of potential interest because, whereas most editing sites in animals occur in noncoding regions associated with repetitive elements, editing in fungi is typically non-synonymous, is under positive selection, and can often lead to changes in coding or STOP sites^52^.

### Down-regulation of *esa-1* impairs the clock

ESA-1 is the catalytic subunit of the NuA4 complex. We were unable to obtain any homokaryotic deletions of the *Neurospora* ortholog of *esa-1* (NCU05218), indicating that it is an essential gene as it is in yeast^16^. To confirm its requirement for normal rhythmicity, a construct bearing a hairpin structure specific for *esa-1* and driven by the *qa*-2 promoter (inducible by QA) was transformed into an ESA-1^V5^ strain, targeting the *csr* locus to create a *ds esa-1* strain affording regulatable knock-down of *esa-1*; circadian rhythms of this strain were monitored using a luciferase assay in the presence or absence of 10^-2^ M QA. In the absence of QA, the *ds esa-1* strain exhibited a robust rhythm similar to that seen in WT (Figure 3a). When 10^−2^ M QA was included in the medium, the robustness of luciferase signals derived from the *frq C-box* was severely impaired (Figure 3a), consistent with ESA-1 being a histone acetyltransferase required for a normal circadian clock. To test whether disruption of ESA-1 can cause a period defect, a dominant-negative mutant of *esa-1* (*esa-1^E395Q^*)^53^ was knocked into a strain bearing *frq*-promoter-driven luciferase at the *his-3* locus. By adding QA, the period in *qa-2-*driven *esa-1^E395Q^* is extended by ∼2 hrs compared with that cultured in the QA-free medium as a control (Figure 3a), suggesting that reduced NuA4 activity can result in a period change; Western blotting confirmed that addition of QA led to a dramatic reduction of ESA-1^V5^ protein only in the *ds esa-1* strain (Figure 3b). To test whether *Neurospora* ESA-1 can acetylate histones H4 and H2A, ESA-1^V5^ was affinity-purified and tested in an *in vitro* acetylation assay using recombinant histones H4 and H2A. ESA-1^V5^ strongly modified histones H4 at lysines 5, 8, 12, and 16 and H2A at lysine 9 (Figure 3c). To directly test whether BRD-8 enhances ESA-1 activity, we used nucleosome core particles (NCPs) as substrate to test the importance of BRD-8 and *in vitro* acetylation assays were carried out with ESA-1^V5^ immuno-precipitated from WT or the Δ*brd-8* background. Surprisingly, we did not see a different activity of ESA-1 in the presence or absence of Δ*brd-8* in the *in vitro* acetylation assay (Figure 3c), which suggests either that Δ*brd-8* might regulate the nucleosomal accessibility of NuA4 *in vivo* and that free histones used in the *in vitro* acetylation assays could not fully recapitulate the chromatin context *in vivo*, or that there is more than one ESA-1-containing complex and that not all ESA-1-containing complexes contain and are regulated by BRD-8.

### *brd-8* is required for normal histone H4 acetylation and Pol II binding at the promoter and **coding region of *frq***

To examine how *brd-8* influences histone H4 and its acetylation status *in vivo*, H4 and acetyl H4 were analyzed by chromatin immunoprecipitation (ChIP) in cultures from the late subjective night. Interestingly, histone H4 densities in Δ*brd-8* increase modestly while the level of acetylated H4 decreases significantly at *frq*, *frh*, *wc-1*, *wc-2*, and other clock-and light-response-associated genes in Δ*brd-8* (Figure 4a and Supplementary Data 2). All these data are consistent with the expected activity of BRD-8, as a NuA4 subunit, in regulating nucleosomal density and histone H4 acetylation genome-wide. Given that BRD-8 interacts with NuA4 subunits, as well as with BYE-1 (Figure 2), a unifying mechanistic model posits that BRD-8 influences the clock through the action of NuA4 and BYE-1 on *frq* transcription. To validate the ChIP-sequencing result of *frq* in Figure 4a, an independent ChIP experiment with new cultures was carried out in the same way as in Figure 4a with the exception of subsequent quantitative PCRs (qPCRs) with *frq*-specific primer sets in place of DNA sequencing. The ChIP-qPCR results agree in general with the ChIP-sequencing data shown in Figure 4a: The pan-histone H4 level at the *frq* gene was increased by approximately two fold, while that of acetylated histone H4 dropped sharply at most sites on *frq* that were examined (Supplementary Figure 6). In a third biological replicate, ChIP-PCR assays were performed to measure the level of pan-and acetyl-histone H4 and Pol II at *frq* (Figure 4b) at a time corresponding to mid-subjective-morning, the peak of *frq* transcription (DD16 in WT and DD19 in Δ*brd-8*). Consistent with results from ChIP-seq (Figure 4a), histone H4 levels at *frq* in Δ*brd-8* were slightly higher than those in WT (Figure 4b), whereas the levels of acetyl histone H4 are significantly lower in Δ*brd-8* than in WT (Figure 4c), and Pol II density within *frq* in Δ*brd-8* is ∼50% than in WT (Figure 4d). The level of WC-1 is lower in Δ*brd-8* (Figure 1) so the reduced levels of histone acetylation and Pol II might be caused by decreased WCC binding to the *frq* promoter. To address this question, ChIP assays were performed using a strain in which *wc-1* driven by the *qa-2* promoter is over-expressed in Δ*brd-8* (Figures 1f and 1g). Interestingly, although WC-1 over-expression in Δ*brd-8* resulted in an even slightly higher binding of WCC to the promoter region of *frq* (Figure 4e) and in lowered histone H4 at *frq* (Figure 4f), the levels of acetylated histone H4 (Figure 4g) and Pol II (Figure 4h) are still much lower than those in WT, suggesting that BRD-8 positively regulates ESA-1 function *in vivo* independent of WCC. ChIP-PCR assays revealed that at DD16, when *frq* transcription is high, BRD-8 associates with both cis-acting elements in the *frq* promoter (*C-box* and *pLRE*) as well as the coding region of *frq* (Figures 4i and 4j). Interestingly, the binding of BRD-8 to the *frq* gene is not dependent on the level of WC-1 expression (Figures 4i and 4j), suggesting that the recruitment of BRD-8 to *frq* does not rely on active transcription.

### Structure-function analysis of the NuA4/BRD-8/BYE-1 complex identifies regions of BRD-8 required for BYE-1 and ESA-1 interaction

To determine whether BRD-8, BYE-1, and NuA4 are in the same or different complexes, BRD-8, BYE-1, and ESA-1 were tagged with V5, 3 x HA, and 3 x FLAG, respectively. Immunoprecipitation with FLAG antibody-conjugated resin was performed, the complex was eluted with 3 x FLAG peptide, and then the ESA-1-3 x FLAG peptide elution was re-immunoprecipitated with V5 antibody. Interestingly, BRD-8, ESA-1, and BYE-1 were seen in the same complex despite a relative low level compared with inputs (Figure 5a), suggesting that they might have a functional overlap. However, in strains lacking BRD-8, BYE-1 no longer interacts with ESA-1 indicating that BRD-8 is required to bridge this interaction and bring BYE-1 into the complex (Figure 5b). To test whether ESA-1 and BRD-8 associate with each other quantitatively, immunodepletion assays were performed using cultures grown in constant light. Immunodepletion of BRD-8^V5^ with V5 antibody only pulled down a fraction of ESA-1 (Figure 5c left), suggesting that different pools of NuA4 complexes exist in the cell. Immunodepletion of ESA-1^3^ ^x^ ^FLAG^ with FLAG antibody pulled down all BRD-8^V5^ molecules (Figure 5c right), suggesting that all BRD-8^V5^ is complexed with NuA4 in the cell. These data explain the results of the *in vitro* histone acetylase assays (Figure 3c), and are consistent with a model in which BRD-8 can modulate the activity of some but not all NuA4 activity in the cell, and may be reminiscent of recent reports that human BRD8 participates in multiple complexes^54^. When *Neurospora* BRD-8^V5^ and BYE-1, ARP-4, or EPL-1 were co-expressed in *E. coli* an interaction was detected by immunoprecipitation (Supplementary Figure 7), indicating that BRD-8 directly associates with NuA4 subunits and BYE-1. BRD-8 is widely conserved among the ascomycete fungi as noted above, indeed more so as compared to a NuA4 subunit described in Saccharomyces but not found in *Neurospora*, EAF5, whose orthologs are restricted to Saccharomyces and closely related species (Supplementary Figure 5). Indeed there is strong evidence suggesting that human BRD8 is the functional homolog of yeast EAF-5 acting as a bridge protein in some complexes^54^.

**Figure 5.**
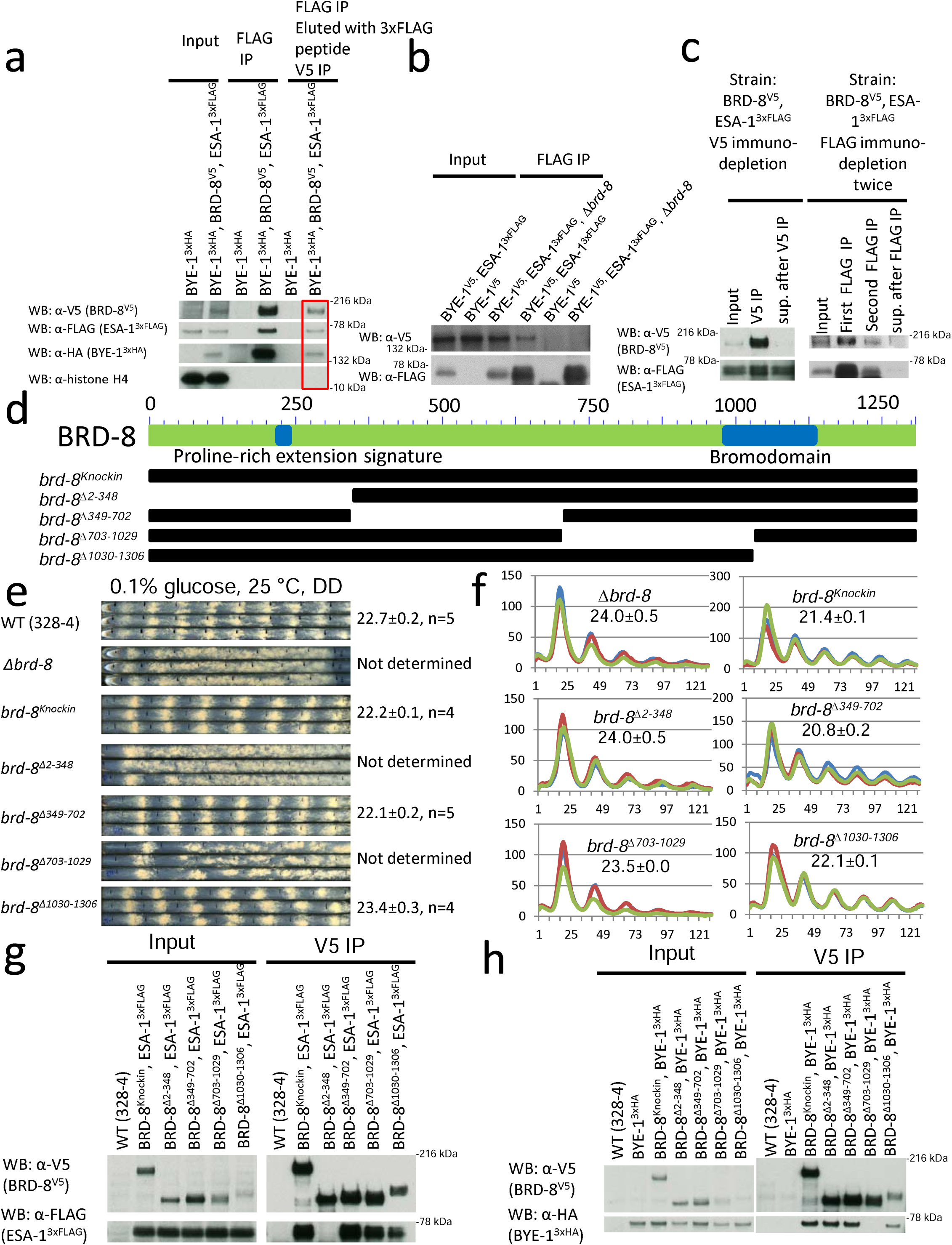
Deletion series of Δ*brd-8* identifies domains responsible for ESA-1 and BYE-1 interactions. Strains indicated were grown in constant light. (**a**) IP showing that fractions of BRD-8, BYE-1, and ESA-1 exist in the same complex. Protein lysate from *bye-1^3^ ^x^ ^HA^/brd-8^V5^/esa-1^3^ ^x^ ^FLAG^* cultured in constant light was first immunoprecipitated with FLAG-antibody resin, washed, and then eluted with 3 x FLAG peptide; for the eluate, a secondary IP was performed using V5-antibody resin; *bye-1^3^ ^x^ ^HA^* serves as the negative control. (**b**) BRD-8 is required for the interaction of ESA-1 and BYE-1. IP with FLAG was performed with three strains of BYE-1^V5^/ESA-1^3^ ^x^ ^FLAG^, BYE-1^V5^, and BYE-1^V5^/ESA-1^3^ ^x^ ^FLAG^/Δ*brd-8* grown in the light. In BYE-1^V5^/ESA-1^3^ ^x^ ^FLAG^/Δ*brd-8*, BYE-1^V5^ was not detected in the “FLAG IP” lane. (**c**) Left, Immunodepletion of BRD-8^V5^ with V5 showing that not all ESA-1 molecules interact with BRD-8; right, immunodepletion of ESA-1^3^ ^x^ ^FLAG^ was done by immunoprecipitating FLAG twice and then the supernatant after the two IPs was pulled down with V5 resin, showing that all BRD-8 molecules associate with ESA-1. (**d**) Schematic representation of domains on BRD-8 and deletion series of *brd-8*. Race tube (**e**) and luciferase (**f**) analyses of *brd-8* deletions were performed as in prior figures (also described in Methods). (**g**) Immunoprecipitations using the deletion series of *brd-8* show that aa 2-348 is required for ESA-1 interaction. V5-IP followed by Western blotting against V5 and FLAG was carried out to check BRD-8^V5^ and ESA-1^3^ ^x^ ^FLAG^ interaction in *brd-8* deletion series. (**h**) Immunoprecipitations using the deletion series of *brd-8* show that aa 703-1029 is required for ESA-1 interaction. *bye-1^3^ ^x^ ^HA^* was targeted at the *csr* locus while BRD-8 mutants were V5-tagged at its native loci. IP by V5 and then Western blotting with V5 and HA as indicated were carried out to check BRD-8^V5^ and BYE-1^3^ ^x^ ^HA^ interaction in *brd-8* deletion series. WT (328-4, an untagged strain) serves as the negative control for IP (**g** and **h**). For Figures 5a–5c, 5g, and 5h, similar results were seen from three independent replicates. Source data are in the Source Data file.

To identify motifs in BRD-8 essential for its circadian function and intra-complex interactions, strains bearing a series of deletions of *brd-8* (Figure 5d) were generated and analyzed by race tube and Western blotting assays. Western blotting analyses showed that of the four *brd-8* deletion strains, only Δ1030-1306 has a reduced protein level (Supplementary Figure 4d). Two of these mutants, missing aa 2-348 and aa 703-1029, showed a conidiation phenotype similar to Δ*brd-8* (Figure 5e), suggesting that these two regions are required for its circadian function. Although analysis of BRD-8’s open reading frame (ORF) revealed a region of bromodomain near the C-terminus of the protein (Figure 5d), deletion of aa 1030-1306 within which the bromodomain is located resulted only in a slightly longer period (23.4 hrs) (Figures 5e and 5f), which suggested that the domain is not essential for its circadian function. This result is consistent with the existence of multiple BRD-8-containing complexes in the cell, only some of which require the bromodomain, and that those requiring the bromodomain are not the functions relevant to circadian timekeeping.

Because ESA-1 and BYE-1 were identified from the BRD-8 interactome analyses and disruption of either gene results in a lengthened circadian period, we checked by immunoprecipitation whether the interaction between BRD-8 and ESA-1 or BYE-1 is influenced in the *brd-8* mutants. Interestingly, BRD-8^Δ2-348^, in which a proline-rich low complexity region is deleted, binds normally to BYE-1 but does not interact with ESA-1 (Figure 5g) while BRD-8^Δ703-1029^ associates with ESA-1 but fails to complex with BYE-1 (Figure 5h), consistent with BRD-8 being the adapter protein for ESA-1 and BYE-1. *brd-8*^Δ*349-702*^ and *brd-8*^Δ*1030-1306*^ showed normal interactions with ESA-1 and BYE-1 (Figures 5g and 5h) and displayed ∼WT circadian phenotypes (Figures 5e and 5f).

### Characterization of roles for BRD-8 in the circadian system

To directly test whether there is a causality between the level of *brd-8* and circadian periods, we made a strain in which the ORF of *brd-8* is under the control of the quinic acid-inducible promoter (*qa-2*) at its native locus. In the presence of QA, the controlled gene is constitutively induced by the *qa*-2 promoter. Interestingly, the level of *brd-8* positively correlated with shortening of the period of the circadian conidiation rhythm within a certain range of QA concentration (10^-5^ to 10^-3^ M) (Figures 6a, 6b, and 6c); however, when the QA concentration reaches 10^-2^ M, the period is slightly lengthened (Figures 6a, 6b and 6c), which might be caused by the indirect effect of the high BRD-8 level on the expression of other proteins that might indirectly influence the circadian clock. The level of BRD-8 is higher when cultured in 2% glucose LCM than in 0.1% LCM, while the ESA-1 level is constant in 2% vs 0.1% glucose LCM (Figure 6d). The BRD-8 level is significantly lower than that of ESA-1 (Figure 6d and Supplementary Figure 8a), consistent with the immunoprecipitation data of Figures 5b and 5c. The BRD-8 protein was highly enriched in the nucleus (Figure 6e), consistent with its role in regulating transcription. BRD-8 does not interact with itself (Supplementary Figure 4e), suggesting that NuA4/BRD-8/BYE-1 does not form an oligomer and BRD-8 might only exist in the NuA4 complex. Because the circadian clock in *Neurospora* globally regulates gene expression, we investigated whether the expression of BRD-8, BYE-1, and several additional NuA4 subunits are also under circadian control. Promoters of *brd-8*, *bye-1*, *eaf-3*, *vid-21*, and *esa-1*, along with *hH2Az*, and *hH4-1* were fused with the codon-optimized LUC gene respectively and transformed to the WT strain at the *csr* locus. The promoters of *brd-8*, *bye-1, eaf-3, vid-21*, and *hH2Az* genes were shown to drive rhythms in LUC expression (Figure 6f), ARP-4 was found to be rhythmic in a separate study^55^, while no obvious circadian rhythmicity was seen for the *esa-1* or *hH4-1* promoters (Supplementary Figure 8b). Furthermore, to better understand and analyze *brd-8* dynamics *in vivo*, we constructed a *brd-8*-luciferase translational fusion reporter that directly tracks BRD-8 protein expression and transformed it into the WT stain. The protein level of BRD-8-LUC oscillated *in vivo* in a circadian manner (Figure 6f), confirming that *brd-8* is a *ccg* (*clock-controlled gene*), and strains bearing BRD-8-LUC show an overt rhythm with a WT period (Figure 6g), indicating that LUC does not impact its function. The data are consistent with a model in which an BRD-8/BYE-1/NuA4 complex controls *frq* expression and thereby the clock, perhaps by modulating transcription elongation, while several subunits of the BRD-8/BYE-1/NuA4 complex are also themselves under circadian control. In this way elements of the BRD-8/BYE-1/NuA4 complex both regulate the clock and are regulated by the clock, contributing to a feedback loop surrounding the core circadian oscillator.

**Figure 6.**
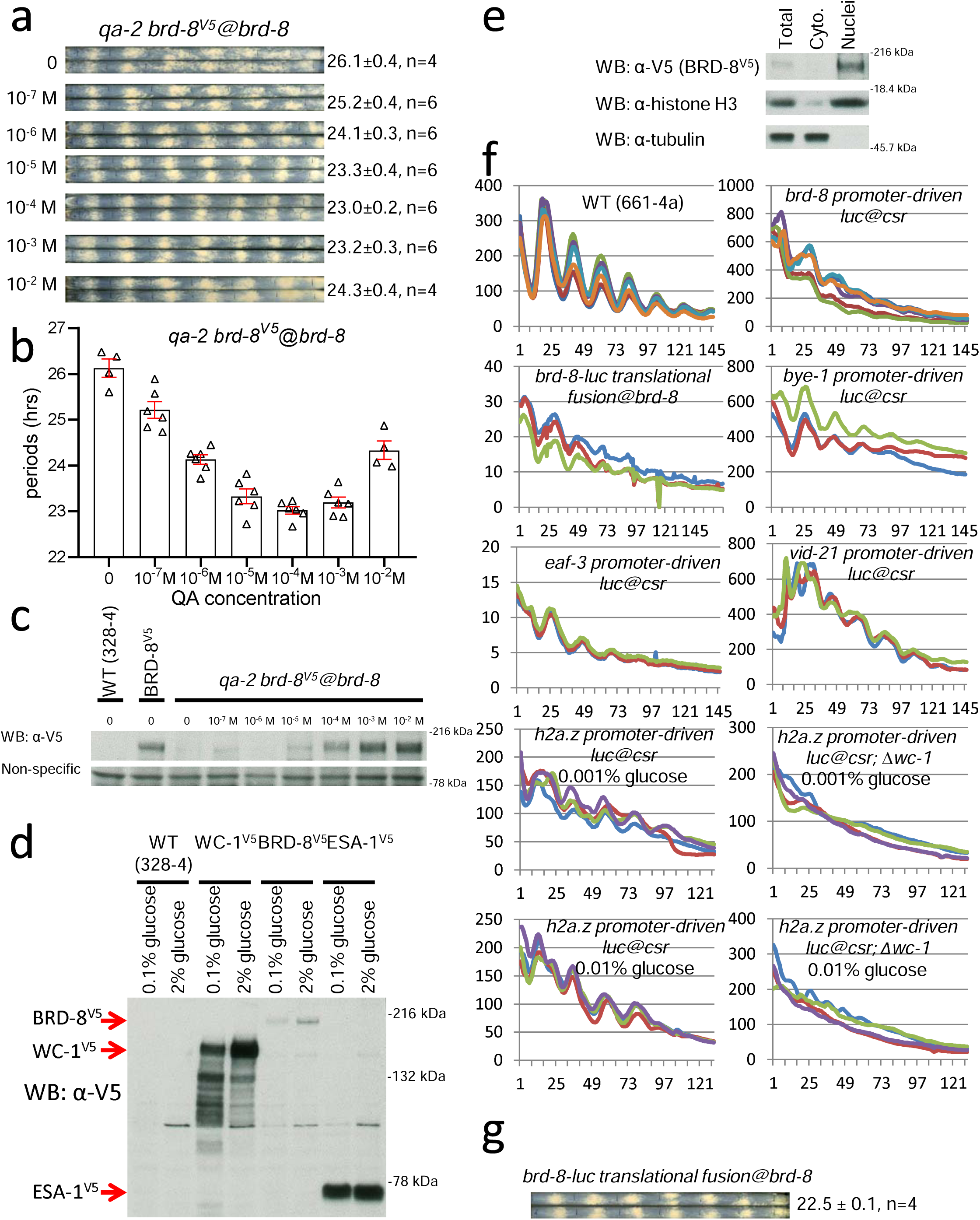
Characterization of *brd-8.* (**a**) Race tube analyses of *qa-2:brd-8^V5^* in the presence of QA at different concentrations as indicated. Strains were cultured on 0.1% glucose race tube medium on race tubes at 25 °C and synchronized by growth in constant light overnight (16-24 hrs) followed by transfer to darkness at the same temperature. (**b**) Plots of periods of *qa-2:brd-8^V5^* in (**a**) Four or six biological replicates of each condition were run and error bars represent standard errors of the means. (**c**) Western blots showing BRD-8 levels in cultures of *qa-2:brd-8^V5^* with different amounts of quinic acid grown in constant light at 25 °C. WT (328-4) and BRD-8^V5^ are the negative and positive controls respectively for the Western blots. (**d**) Western blotting of *brd-8^V5^*, *wc-1^V5^*, and *esa-1^V5^* in cultures grown in different amount of glucose (0.1 or 2%) in constant light. V5 was inserted to the C-termini of BRD-8, WC-1, and ESA-1 at their native loci respectively, and Western blotting against the same tag was used to fairly compare their protein levels. Red arrows point to BRD-8^V5^, WC-1^V5^, and ESA-1^V5^ as indicated. (**e**) Nuclear fractionation of BRD-8^V5^ from a culture grown in constant light. The nuclear fractionation was performed as previously described^90^; histone H3 and tubulin were followed as a nuclear and cytoplasm marker respectively. For Figures 6c–e, similar results were observed from three replicates. (**f**) Bioluminescent analyses of luciferase expression under the control of promoters of *brd-8*, *bye-1*, *eaf-3*, *vid-21*, or *h2a.z* individually 25 °C; LUC fused to the C terminus of BRD-8 ORF (translational fusion at the locus of *brd-8*) was also measured by the bioluminescent analysis at 25°C. (**g**) Race tube analysis of *brd-8*-LUC with 0.1% glucose race tube medium in the dark at 25 °C. Source data were put in the Source Data file.

### Inhibition of CDK-9 further lengthens the circadian period of **Δ***brd-8*

If mutations in the NuA4/BRD-8/BYE-1 complex impact the clock by affecting transcription elongation, then other mutations or chemicals impacting elongation should have similar effects on the clock and, importantly, loss of BRD-8 might be expected to sensitize the clock to treatments affecting elongation. Phosphorylation of Ser2 in the Pol II C-terminal domain (CTD) is required for Pol II elongation processivity and CDK-9 (SGV1 in Saccharomyces) is the kinase phosphorylating Ser2. To test whether inhibition of CDK-9 causes a period effect on Δ*brd-8,* strains were grown on medium containing different concentrations of AZD4573^56–59^, a highly specific CDK-9 inhibitor, and circadian oscillations were monitored by luciferase assays. A clear trend of period lengthening effects was observed in WT in the presence of 1-100 μM AZD4573 while Δ*brd-8* shows even a stronger dose effect on period changes than WT, especially 100 μM at which concentration Δ*brd-8* becomes completely arrhythmic while WT is ∼ 6 hr longer than seen under no drug treatment (Figure 7a). CDK-9 was also downregulated independently by replacing its native promoter with the *qa-2* promoter; luciferase assays were conducted in the absence of QA. Compared with Δ*brd-8* (Figure 7a), Δ*brd-8, qa-2:cdk-9* displays a further lengthened period by 2-3 hrs (Figure 7b). The data suggest that the absence of *brd-8* makes Pol II more susceptible to inhibition and both chromatin architecture and robust Pol II elongation are required for the period length determination.

**Figure 7.**
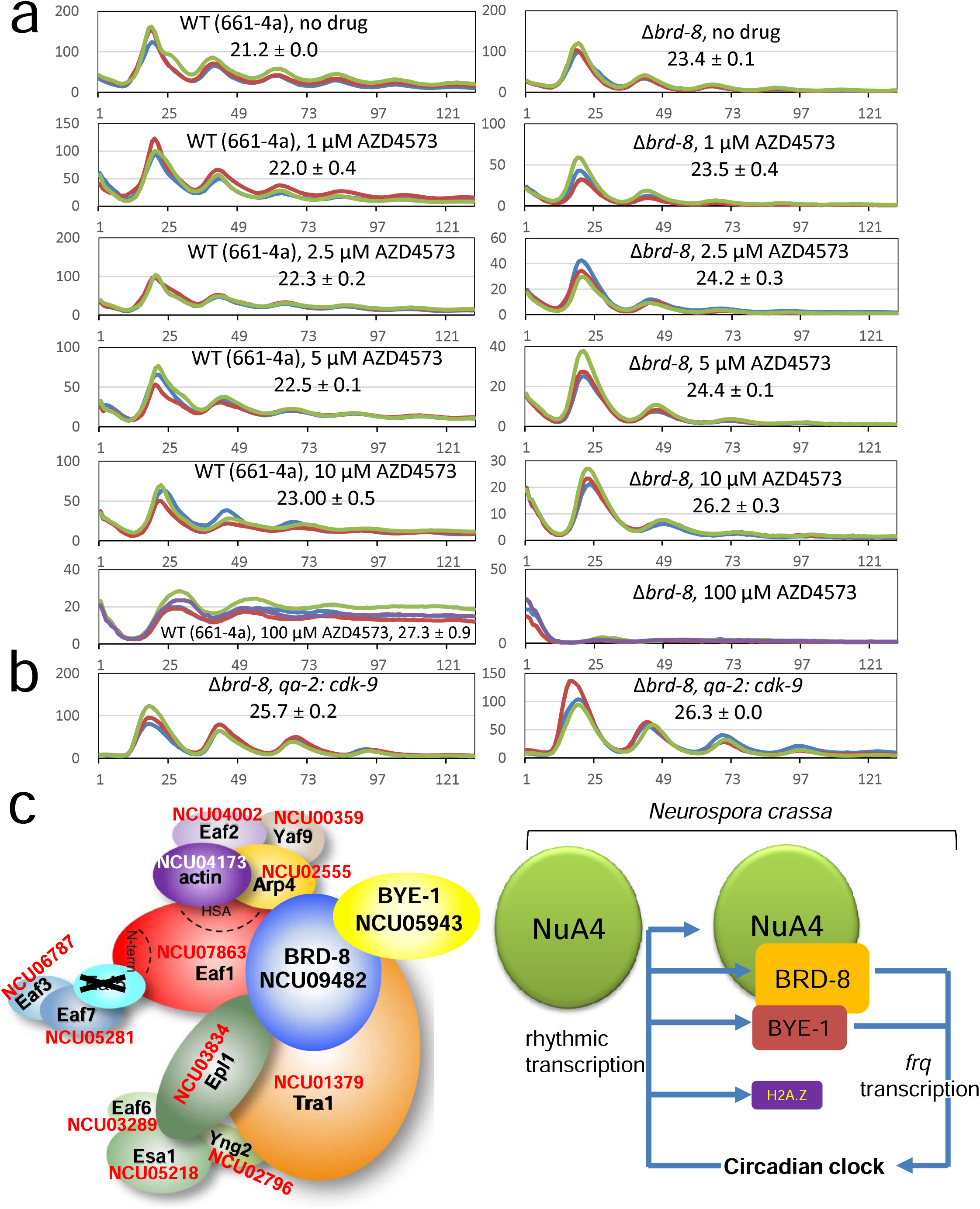
Impairment of transcription elongation lengthens circadian period, leading to a model in which BRD-8 and BYE-1, in complex with NuA4 subunits, maintain circadian period length. (**a**) Pharmacological inhibition of CDK-9 results in period lengthening and further lengthens the circadian period of Δ*brd-8*. Bioluminescent analyses of WT and Δ*brd-8* in the presence or absence of AZD4573, an inhibitor of CDK-9, at concentrations of 0, 1, 2.5, 5, 10 and 100 μM as indicated in the figure. AZD4573 was added to 0.1% glucose race tube medium in a 96 well plate to the final concentrations indicated, and strains inoculated and grown in the light at 25 °C overnight were transferred to the dark at the same temperature to record bioluminescent signals. (**b**) Genetic downregulation of CDK-9 results in period lengthening. Bioluminescent analyses of two independent Δ*brd-8* strains in which the promoter of *cdk-9* was replaced by the *qa-2* promoter to down-regulate expression of *cdk-9* when QA was absent in the medium. (**c**) Working model. On the left is the conceptual model of the NuA4 complex described here, modified from previous work^95^ to add auxiliary subunits BRD-8 and BYE-1 and to show interactions established here along with the corresponding *Neurospora* gene numbers; Eaf5 is crossed out to reflect its absence (Supplementary Figure 5). Right: there are at least two types of NuA4 complexes in *Neurospora* with or without BRD-8 and BYE-1. BRD-8 and BYE-1 are required for normal *frq* transcription and circadian period determination; transcription of *brd-8*, *bye-1*, *eaf-3*, *vid-21*, and *h2a.z* are under circadian control.

## DISCUSSION

The *frequency* gene encodes the central scaffold protein that organizes the negative arm repressive complex in the *Neurospora* clock, playing the same role in *Neurospora* as that ascribed to the PER proteins in the mammalian clock; as a result, its expression is carefully regulated. *frq* is rhythmically transcribed and translated in the dark^10^, a result of chromatin remodelers and modifiers that co-execute their effects at the *frq* locus, and most research on its transcriptional regulation has focused on transcription initiation at the *C-box* rather than on steps in transcription elongation. During daylight the *frq* gene lies in a region largely free of nucleosomes and showing rapid nucleosome turnover^11^, and in the dark several distinct chromatin remodeling complexes have been identified in controlling *frq* expression, including CSW (Clockswitch)^60^, CHD1 (chromodomain helicase DNA–binding)^61^, SWI/SNF (SWItch/Sucrose NonFermentable)^62^, CLOCK ATPase (CATP)^63^, Centromere Binding Factor 1 (CBF-1)^64^, and IEC-1-INO80 complex^65^. Histone modifications can also lead to changes in gene expression, especially methylation and acetylation of N-terminal tails of histones which are known to influence condensation as well as serving to call down modifiers^66–69^; indeed one study has reported that the lysine methyltransferase SET1 is required for methylation of lysine 4 of histone H3 and normal circadian expression of *frq*^70^. Here, by a circuitous route beginning with an unbiased genetic screen, we have identified a phylogenetically conserved auxiliary subunit of the canonical NuA4 histone acetylase complex, shown that it also complexes with elements involved in transcription elongation, and that in the aggregate mutations in these elements impact *frq* expression and the progress of the circadian clock. In the process we also discovered that there are multiple NuA4 complexes, presumably with altered activities, and that some include an ortholog of the regulator of transcription elongation, BYE-1, first described in Saccharomyces^24^ but with orthologs in humans^26^. Genetic and pharmacological data are consistent with a model in which the NuA4 complex that includes BRD-8 and BYE-1, but not the canonical complex, is involved in controlling *frq* expression in the service of the clock (Figure 7c). Given that the expression of a number of NuA4 subunits is in turn regulated by the clock, it seems likely that events surrounding histone acetylation and transcription elongation describe a feedback loop surrounding the core clock.

This work follows the lines of Belden, He, Liu, and others^60–64^ showing that normal transcription of *frq* is required for normal clock function, and also the thorough study of Petkau et al.^23^ which revealed a role for Pol II pause/release and transcription elongation in the mammalian clock. In this way analysis of circadian rhythmicity provides an especially sensitive tool for dissection of transcriptional regulators because differences in period length are biologically meaningful, more nuanced than life/death (e.g. identification of essential genes) and can be reliably measured with precision of +/-10%. The BRD-8-containing NuA4 complex however, like the other chromatin-regulating and clock-affecting complexes, would not be considered a core clock component because its role in rhythmicity is solely conferred by its action on *frq* expression whose rhythmicity is driven through WCC, and the rhythm persists in the absence of BRD-8; instead these are all clock-affecting. This said, the data do show that not all genes are bound by nor their expression influenced by complexes that contain BRD-8 and BYE-1. Instead action of these NuA4-associated factors is restricted, although they do appear to be quite important when present: Data shown in Figures 5c and 6d suggest that BRD-8 only accounts for a small fraction of ESA-1-containing NuA4 complexes, but deletion of *brd-8* resulted in ∼50% reduction of histone H4 acetylation at the *frq* locus, suggesting a higher activity of the BRD-8-containing NuA4 complex.

### Composition of NuA4 complexes

In this study several new subunits associating with the NuA4 complex were identified including BRD-8 and BYE-1 (Figure 7c). In yeast, the NuA4 complex is comprised of 13 subunits (Figures 2c and 7c). In *Neurospora*, however, no gene having homology to *eaf-5* exists, and indeed orthologs of EAF5 are found only in Saccharomyces and closely related species (Supplementary Figure 5). Among the aspects of physiology possessed by fungi outside of the Saccharomycotina are responses to environmental stimuli including light and time-of-day, circadian biology. This correlation suggests the possibility that multiple NuA4 complexes with more subunits may have evolved to support additional aspects of physiology.

Another aspect of regulation found in *Neurospora* but not Saccharomyces is RNA A-to-I editing. In the fungi this mainly occurs in coding regions and results in nonsynonymous changes to proteins encoded^51^, which dramatically extends the proteomic complexity during translation and thus can broadly impact cellular and physiological events. Interestingly, mRNAs of all NuA4 subunits undergo multiple A-to-I editing events in the sexual phase (Figure 2b), which may enable histone H4 acetylation to epigenetically control the chromatin state favoring sexual development and adaptation by influencing chromatin events such as gene expression, DNA repair, and genome duplication.

### Dual regulation between circadian clock and basic transcriptional processes

BRD-8 influences rhythmicity through chromatin events. In Δ*brd-8*, nucleosome density is higher at *frq* while histone H4 is less acetylated than in WT, both of which effects will impede *frq* transcription. *frq* is known to lie in a region characterized by very rapid nucleosome turnover^11^ so in affecting nucleosome density and modification, loss of *brd-8* might be expected to cause reduced *frq* expression and a long period which can be rescued by adding a second copy of the *frq* gene to the Δ*brd-8* strain. The data are consistent with models in which timely expression of *frq* is required for bringing kinases to phosphorylate WCC and effect repression in the circadian cycle. As FRQ and WC-1 are present in similar amounts and FRQ-promoted WCC phosphorylation is a slow process, at least in part because FRQ is predominantly cytoplasmic^9^, the level of FRQ needs to be sufficient to bring about repression to WCC.

BYE1 was originally identified in Saccharomyces through a mutation that bypassed the requirement for the essential prolyl isomerase ESS1^24^. Ess1-induced conformational changes are seen as attenuating Pol II elongation, thus helping to coordinate the assembly of protein complexes on the Pol II CTD and thereby regulating transitions between multiple steps of transcription^24^. BYE1 binds directly to Pol II during early transcription elongation and tethers surrounding histones containing active PTMs, perhaps to prevent loss of histones during polymerase passage through chromatin^26^ but also slowing transcription. Although in Saccharomyces BYE1 works as a free protein, its action(s) in *Neurospora* in conjunction with NuA4 may be similar to those seen in yeast, acting on elongation through regulating Pol II CTD or just enhancing ESA-1 activity on chromatin, an activity not reported in Saccharomyces. In either case, a seeming paradox in the circadian data related to loss of BRD-8 and BYE-1 arises from the facts that (1) loss of BRD-8 which results in increased nucleosome density is expected to slow transcription but (2) loss of BYE-1, a negative regulator of transcription elongation^26^, would be expected to increase transcription elongation; yet both result in period lengthening. One clue to this lies in the data from Figure 5 showing that BRD-8 forms the bridge between ESA-1 and BYE-1; hence, loss of BRD-8 results in parallel loss of BYE-1 from the NuA4 complex. This still leaves unexplained, however, how single loss of BYE-1 in the presence of normal BRD-8 results in modest period lengthening. A possible resolution to this lies in the data of Figure 6 which detail the dosage effects of BRD-8 on rhythmicity. While complete loss of BRD-8 slows period to the greatest extent, the highest expression of BRD-8 also results in modest period lengthening, comparable to loss of BYE-1. It may be that overexpression BRD-8 has consequences on transcription similar to loss of BYE-1, and that normal rhythmicity requires a happy medium of regulators of transcription elongation, allowing it to proceed but not to proceed so quickly as to interfere with ordered assembly of protein complexes on the Pol II CTD.

### BRD-8, the NuA4 complex, and roles for acetylation

The unexpected identification of a human BRD8-like protein associated with a fungal NuA4 acetyltransferase complex, and the role of acetylation in the fungal clock, warrant comment. In mammals, Brd8 is expressed in multiple isoforms, some with one and some with two bromodomains, although expression of the two-bromodomain isoform is restricted to testes whereas the single bromodomain isoform is ubiquitously expressed (e.g., GTExportal.org) and it is similar in size to *Neurospora* BRD-8 that likewise has a single bromodomain. While as noted above, BRD8 is associated with multiple mammalian complexes including NuA4/TIP60, the association of a bromodomain-containing protein in a fungal NuA4 complex has been debated (reviewed in^49,71^). Bdf1 in the *Saccharomyces cerevisiae* SWR1 complex, has a single bromodomain as does Bdc1, a component of the NuA4 histone acetyltransferase complex In *Saccharomyces pombe* (reviewed in^49,71^). However, the overall amino acid identity of *Neurospora* BRD-8 with Bdf1, and Bdc1 is relatively low (6.96%, and 3.9%, respectively, similar to background), Bdf1 and Bdc1 are a third of the size of BRD-8, and perhaps most significantly, neither Bdf1 nor Bdc1 are BLASTP hits with BRD-8. Bdf1 is, however, a component of the yeast SWR complex which contains additional components orthologous to mammalian NuA4/TIP60 components, prompting the observation that the human NuA4 complex corresponds to a near-perfect fusion of the two distinct complexes in yeast, namely NuA4 and SWR1 (Doyon and Cote, 2004). This prompted us to re-examine the mass spectrometry data (Figure 2 and Supplementary Data 1) for presence of putative SWR complex components. No peptides from subunits exclusively in SWR (i.e. orthologs of Bdf1, Vps72, Rvb1, Rvb2) were recovered from the BRD-8 V5 purification (Supplementary Data 1), suggesting that the presence of BRD-8 in association with NuA4 is more like that seen in mammals than the association of bromodomain-containing proteins sometimes seen in yeasts.

A second point of interest sparked by the identification of BRD-8 in association with NuA4 is the potential role of acetylation in rhythmicity. This question is nicely set up by the elegant study of Petkau et al who showed in mammals that the NuA4/TIP60 complex acetylates BMAL1 (the ortholog of *Neurospora* WC-1), providing a docking site that brings the BRD4-P-TEFb complex to DNA-bound BMAL1, promoting release of Pol II and elongation of circadian transcripts and revealing a role for control of transcriptional elongation in the mammalian clock^23^. At first glimpse there are clear parallels to the BRD-8 story here, in particular because WC-1 has been reported to be acetylated^72^; however, the similarities weaken when examined more closely. *Neurospora* does have a Brd4 ortholog (BDP-3 encoded by NCU08423) and, unlike BRD-8, it is essential as is Brd4 in humans. The acetylation of WC-1 appears only to be important for light-regulation^72^, and perhaps most importantly, the structure/function data from Figure 5d–h plainly indicate that the bromodomain of BRD-8 is not required for its circadian function. The mass spectrometry data from Supplementary Data 1 suggest that BRD-8 may interact with a number of different proteins and complexes, and if so it is like human Brd8 which interacts with at least 135 proteins (https://thebiogrid.org/116108), is ubiquitously expressed, and has roles in a host of processes; in both systems it seems plausible that not all of these roles would require the bromodomain. In the present case, BRD-8 uses its unstructured region to interact with BYE-1, and deleting its single bromodomain still results in a normal period while removal of two disordered domains individually phenocopies the whole gene knockout of *brd-*8 (Figures 5d–h). Although BRD-8 provides no evidence for a role of bromodomain-mediated acetylated residue binding, a more remote route to achieve this must be mentioned. The human ortholog of BYE-1 is DIDO1 (death inducer-obliterator 1) which has been shown in a high throughput affinity capture-ms screen^73^ to interact with Brd4, the protein shown by Petkau et al^23^ to mediate the pause/release of Pol II from DNA-bound BMAL1. If the same mechanism worked in *Neurospora*, it should be mediated by the *Neurospora* ortholog of Brd4 (BDP-3) and be independent of the BRD-8/BYE-1 association, and if so the period lengthening seen from loss of either BRD-8 or BYE-1 alone should be additive in the double mutant Δ*brd-8, Δbye-1;* however, this is not the case (Figures 1a, and 2d). Additionally, there is no evidence (mass spectrometry in Supplementary Data 1 or targeted immunoprecipitation of BRD-8 followed by blotting for FRQ or WCC) for interaction of BRD-8 with FRQ or WCC. Lastly BDP-3 (the *Neurospora* ortholog of human BRD4) does not appear in the mass spectrometry interactome data (Supplementary Data 1) indicating that even if BYE-1 has a circadian role through BDP-3, it cannot be linked in any mechanistic way to BRD-8.

A wealth of research in recent years has established mechanisms through which epigenetic regulation plays an important role in controlling circadian systems (reviewed in^74–78^); however, the reverse has rarely been confirmed (reviewed in^79^). Here we have shown that expression of *hH2Az*, as well as a number of NuA4 subunits including *brd-8* gene and protein are circadianly controlled (Figure 7c), linking the circadian clock to the basic cellular machinery. This prompted us to examine online databases for circadianly regulated genes in mammals (CIRCA: Circadian gene expression profiles; http://circadb.hogeneschlab.org) for evidence of circadian regulation of mammalian NuA4/TIP60 components. Interestingly, strong albeit sometimes low amplitude rhythms were seen for gene expression of *brd8* as well as *Trrap, RUVBL1, and DMAP1*. This suggests that in mammals, as in *Neurospora*, the events surrounding histone acetylation and transcription elongation describe a feedback loop surrounding the core clock that serves to amplify and stabilize the rhythm, contributing to persistence. These data also provide evidence for a mechanism through which the circadian clock can control gene expression through an indirect mechanism not directly mediated by the WCC or CLOCK/BMAL1 complex, the principal means of circadian output control.

## Methods

### Strains and growth conditions

328-4 (*ras-1^bd^, A*) was used as a clock-WT strain in the race tube analyses and 661-4a (*ras-1^bd^, A, his-3::C-box-driven luc*), WT in the luciferase assay; this contains the *frq C-box* fused to codon-optimized firefly *luciferase* (transcriptional fusion) at the *his-3* locus. *Neurospora* transformation was performed as previously reported^62^. To test the rhythmicity of gene expression, promoter-driven *luciferase* (*LUC*) reporter constructs were generated by transforming four PCR products [5’ of *csr*, promoter region of gene, codon optimized *luciferase* sequence, and 3’ of *csr*] with digested plasmid (*pRS426*) into yeast ^80^. Race tube medium contains 1 x Vogel’s salts, 0.17% arginine, 1.5% bacto-agar, and 50 ng/mL biotin with glucose at 0.1%, and liquid culture medium (LCM) contains 1 x Vogel’s, 0.5% arginine, 50 ng/mL biotin, and 2% glucose. quinic acid (QA) was added into race tube medium with certain concentrations as indicated. Unless otherwise specified, race tubes were cultured in constant light for 16–24 hrs at 25 °C and then transferred to the dark at 25 °C^81^. BRD-8, BYE-1, and NuA4 deletion strains generated by the *Neurospora* genome project were obtained from Fungal Genetics Stock Center (FGSC).

### Protein lysate and Western blot

Procedures for preparation of protein lysates and Western blots (WB) were followed as described^8,82^ For WB, 15 µgs of whole-cell protein lysate were loaded per lane in a 3-8% tris-acetate or 4-12 bis-tris SDS gel. Antibodies against WC-1, WC-2, FRQ, and FRH have been described previously. V5 antibody (Thermo Pierce), FLAG antibody (Sigma-Aldrich), and HA (Abcam, ab9110) were diluted 1:5,000 for use as the primary antibody. All uncropped and unprocessed whole blots used for figure preparations were deposited into the Source Data file except for the ones for Supplementary Figure 4a, which were damaged permanently in a lab fire.

### Identification of BRD-8 interactors

VHF-tagged BRD-8 were purified with the same method applied for isolation of C-terminal VHF-tagged WC-1^8,62^. 30 g of *Neurospora* tissue (*brd-8^VHF^*) was used for purification of BRD-8. To exclude the possibility that associations with chromatin impacted identification of BRD-8-interactors in purifications, we (1) vortexed lysate at the top speed to mechanically disrupt chromatin: 10s on/10s off (put on ice) for a total of 2 min; (2) extensively sonicated protein lysate; (3) included benzonase nuclease In the protein lysis buffer; benzonase is widely used for the removal of nucleic acids (both DNAs and RNAs) from protein samples^83–86^. To recover potential weak interactors, 5 g of tissue from *brd-8^V5^* were purified with V5 antibody (Thermo Pierce)-conjugated Dynabeads (Life Technologies). A single protein band or the whole lane^87^ were cut off from the gel as indicated in the figure and used for mass spectrometry.

### Mass spectrometry analysis

Affinity purification of BRD-8’s interactors using VHF-tagged BRD-8 (the left gel in Figure 2a) was performed twice (two biological replicates) with similar results, and identification of proteins from excised bands was carried out once with mass spectrometry. Interactor identification from BRD-8^V5^ (the middle and right gels in Figure 2a) was performed three times (three biological replicates), similar band profiles were observed, and specific bands or whole gel lanes were analyzed once by mass spectrometry. Validations for interaction of BRD-8 with its interactors identified from mass spectrometry analyses were performed with strains bearing epitope-tagged target proteins, and results were shown in Figure 2c. One negative control used in the BRD-8’s interactor identification was an untagged *Neurospora* strain (328-4 [*ras-1^bd^*, A]). Six samples were analyzed by mass spectrometry.

Samples were separated by SDS-PAGE gel electrophoresis and Coomassie-stained. Gel bands or lanes were excised, destained, and digested with trypsin in 50 mM Ammonium bicarbonate overnight at 37°C. Peptides were extracted using 5% formic acid / 50% acetonitrile (ACN), desalted, and dried. Peptides were analyzed on a Fusion Orbitrap mass spectrometer (Thermo Scientific) equipped with an Easy-nLC 1000 (Thermo Scientific).

Peptides were resuspended in 5% methanol / 1% formic acid and loaded on to an analytical resolving column (35 cm length, 100 μm inner diameter, ReproSil, C18 AQ 3 μm 120 Å pore) pulled in-house (Sutter P-2000, Sutter Instruments, San Francisco, CA) with a 45-min gradient of 5–34% LC-MS buffer B (LC-MS buffer A: 0.0625% formic acid, 3% ACN; LC-MS buffer B: 0.0625% formic acid, 95% ACN). The Fusion Orbitrap was set to perform an Orbitrap MS1 scan (R = 120K; AGC target = 2.5e5) from 350 to 1,500 m/z, followed by HCD MS2 spectra detected by Orbitrap scanning (R = 15K; AGC target = 0.5e5; max ion time = 50 ms) for 2 sec before repeating this cycle. Precursor ions were isolated for HCD by quadrupole isolation at width = 0.8 m/z and HCD fragmentation at 29 normalized collision energy (NCE). Charge state 2, 3, and 4 ions were selected for MS2. Precursor ions were added to a dynamic exclusion list ± 10m ppm for 15 s.

Raw data were searched using COMET^88^ (release version 2014.01) in high resolution mode against a target-decoy (reversed)^89^ version of the *Neurospora* proteome sequence database (UniProt; downloaded 3/2017) with a precursor mass tolerance of +/-1 Da and a fragment ion mass tolerance of 0.02 Da, and requiring fully tryptic peptides (K, R; not preceding P) with up to three mis-cleavages. Static modifications included carbamidomethylcysteine and variable modifications included: oxidized methionine. Searches were filtered using orthogonal measures including mass measurement accuracy (+/-3 ppm), Xcorr for charges from +2 through +4, and dCn targeting a <1% FDR at the peptide level.

The list of identified interactors (Supplementary Data 1) was searched manually for proteins whose peptides were specifically enriched in the sample of BRD-8^V5^ but not in the negative control (328-4, untagged). In the search for BRD-8^V5^’s interactors, no statistical tests were taken, because we worried that this would rule out certain real targets due to their potential digestion or/and coverage issues in mass spectrometry analyses. Instead, we validated the identified interactors individually by immunoprecipitation assays with epitope-tagged proteins (Figure 2c).

### Immunoprecipitation (IP)

IP was performed as previously described^90,91^. Briefly, 2 mgs of total protein were incubated with 20 μL of V5 agarose (Sigma-Aldrich, Catalog # 7345) or FLAG M2 resin (Sigma-Aldrich, Catalog # A2220) as indicated rotating at 4 °C for 2 hrs. The agarose beads were then washed twice with the protein extraction buffer (50 mM HEPES [pH 7.4], 137 mM NaCl, 10% glycerol, 0.4% NP-40) and eluted with 50 µL of 5 × SDS sample buffer at 99 °C for 5 min.

### Chromatin immunoprecipitation (ChIP)

ChIP experiments were done as previously described using fresh tissues^62^. Primer sets against *C-box* and *pLRE* were the same as in a previous publication^8^. Primer sets against the *frq* locus used in Figure 4a were derived from a previous publication^92^. “A” in the first “ATG” of the full-length *frq* coding region is designated as “+1” and the upstream nucleotides are labelled “-“ accordingly (without “0”). *frq C-box* is from –2,878 to –2,624 nt, the amplicon with the *frq1* primer set is from –2,754 to –2,644 nt while that by the *C-box* primer set is from –2,790 to –2,678 nt. ChIP using *Neurospora* tissue: *Neurospora* tissue was cross-linked with 3% formaldehyde for 15 min and quenched with 0.25 M glycine for 5 min. Tissue was then washed with PBS three times and vacuum-dried; 0.4 g was weighed, cut into 9 pieces, and each piece soaked in 0.5 ml SDS lysis buffer (50 mM Tris/HCL [pH 8.0], 1% SDS, 5 mM EDTA) containing Roche protease inhibitors (Sigma-Aldrich, Catalog # 11836170001). The soaked tissue was first sonicated for 8 sec for three times at 30% amplitude with a Bronson sonicator equipped with a microtip, and then further sonicated in a water bath sonicator 5 times for 5 min each with an interval of 30 sec on and 30 sec off. The sonicated cell lysate was cleared of cellular debris by an 8,000 rpm centrifugation twice for 5 min. 200 μl supernatant was saved and added to 1.8 ml RIPA buffer; 15 μL protein A magnetic beads and 2 μL V5 antibody (Abcam, Catalog # ab9116) were added to the mixture followed by rotation at 4 °C overnight. The following morning, immunoprecipitated DNA was washed with buffer A-D (buffer A: 20 mM Tris/HCL [pH 8.0], 0.1% SDS, 2 mM EDTA, 1% TritonX-100; buffer B: 20 mM Tris/HCL [pH 8.0], 0.1% SDS, 2 mM EDTA, 1% TritonX-100, 500 mM NaCl; buffer C: 10 mM Tris/HCL [pH 8.0], 0.25 M LiCl, 1 mM EDTA, 1% TritonX-100, 1% sodium deoxycholate; buffer D: 10 mM Tris/HCL [pH 8.0], 1 mM EDTA) once each, eluted with Elution buffer (0.1 M NaHCO_3_, 1% SDS), reverse crosslinked with NaCl at the final concentration of 0.2 M, and purified with Qiagen PCR purification kit.

### Ion ChIP-Seq Library Preparation

The purified ChIP DNA was end-repaired and ligated to Ion-compatible barcode adapters using Ion XpressTM Plus Fragment Kit (Catalog # 4471269) and Ion XpressTM Barcode Adapters (Catalog # 4471250). The final libraries were purified with two rounds of AMPure XP Bead capture to size select fragments between 160 to 340 bps in length. The emulsion clonal bead amplification to generate bead templates for the Ion Torrent platform was performed on the Ion Chef System (Thermo Fisher Scientific) with the Ion PI™ Hi-Q™ Chef Kit (Thermo Fisher Scientific, Catalog # A27198) and Ion PI™ Chip Kit v3 (Thermo Fisher Scientific, Catalog # A26771). Sequencing was done using the Ion PI™ Hi-Q™ Sequencing 200 Kit (Thermo Fisher Scientific, Catalog # A26772) on Ion Proton sequencer with sequencing data processing using the Torrent Suite TM Software (Ver. 4.0.2) on the Torrent server.

Samples were mapped to the *Neurospora crassa* genome (from GenBank) using TMAP (Ion Software, https://github.com/iontorrent/TMAP). Mapping efficiency and read quality were visualized using htseq-qa (http://htseq.readthedocs.io/en/release_0.9.1/). Peaks were called using MACS2 (https://github.com/taoliu/MACS), and peaks were annotated with ChIPseeker (Bioconductor, https://bioconductor.org/packages/release/bioc/html/ChIPseeker.html). In addition to MACS2, the SICER software (https://home.gwu.edu/~wpeng/Software.htm) was used to identify peaks and identify differentially abundant peaks between WT and mutant strains. For comparison of WT vs. control in histone and acetylated histone, three runs apiece were performed with window sizes of 200, 400 and 600, and gap sizes of 600, 1,200 and 1,800, respectively. Redundant peaks between the three runs (within 2,500 bp) were eliminated with a simple R script. Another script was used to identify genes within 1,000 bp from either border of the peaks.

### Histone acetylation assay *in vitro*

The histone acetylation assay was performed as previously described with modifications^93^. Briefly, ESA-1 was immunoprecipitated from the *esa-1^V5^* strain as described in the section of “IP” with V5 antibody-conjugated resins. Immunoprecipitated-ESA-1^V5^-bound resins were washed twice with phosphate buffered saline containing 0.1% Tween 20 and once with the acetyl-transferase assay buffer (50 mM Tris-Cl pH 8.0, 10% glycerol, 10 mM butyric acid, 0.1 mM EDTA, 1 mM DTT, 1 mM PMSF). 50 μL of the acetyl-transferase assay buffer containing 10 μM acetyl CoA (Sigma-Aldrich) was added to the washed resins. Recombinant human Histone H2A (New England BioLabs, Catalog # M2502S) or H4 (New England BioLabs, Catalog # M2504S) respectively was added to the mixture at a final concentration of 0.1 mg/mL. Acetylation reactions were incubated at 30 °C for 1 hr, followed by adding 50 μL of 5 x SDS-PAGE sample buffer, heated at 99 °C for 5 min, electrophoresed in a 4–12% bis-tris SDS-PAGE gel, and transferred to a PVDF membrane. Acetylated histone H4 and H2A were detected by immunoblotting with rabbit anti-acH4 (UBI, Catalog # 06–866, 1:1000 dilution) and anti-acH2A K9 (Millipore Sigma, Catalog # 07-289), respectively.

### RT-quantitative PCR

*Neurospora* mRNA was isolated with TRIzol reagent (Life Technologies) and cDNA synthesis was done using SuperScript III first strand synthesis kit (Life Technologies) with 3 µg of purified RNA^47^. Real-time PCR was performed with QuantiTect SYBR green RT-PCR kit (Qiagen) in an ABI 7500 Fast system. Primers against *frq* and *wc*-1 genes were from a previous publications^47,94^. Normalization for RT-qPCR was to NCU08964, a gene identified previously as among the most constantly expressed under different conditions and times or day, and least responsive to changes in media^40^.

### Other techniques

Mass Spectrometry was performed as previously described^62^. Luciferase assays were performed as previously described^8,81^. If QA was used in the luciferase assay, it was indicated in the figures. Nuclear preparation was performed as reported^90^.

### Data availability

The mass spectrometry and ChIP-seq data generated in this study have been deposited in the MassIVE database under accession: MSV000089415 (link: https://massive.ucsd.edu/ProteoSAFe/dataset.jsp?task=74b7985c9ee9409485fb212a918c6b4b, PDX#: PXD033576) and NCBI SRA under accession: PRJNA834768 (https://www.ncbi.nlm.nih.gov/sra/PRJNA834768), respectively. Source data are provided with this paper.

## Acknowledgments

We thank the Fungal Genetics Stock Center at Kansas State University for *Neurospora* strains, Yi Liu and Xiao Liu at The University of Texas Southwestern Medical Center for help with transforming the ds RNA strain, and Christina Kelliher and Wei Wang in the Dunlap and Loros labs for technical assistance in visualizing the ChIP-seq data and conducting menadione race tube assays respectively. We are also grateful to Mark E. Adamo in the Kettenbach lab for help in depositing the raw mass-spectrometry data in the MassIVE database. This work was supported by grants from the National Institutes of Health to J.C.D. (R35GM118021), J.J.L. (R35GM118022), A.N.K. (R35GM119455), and Department of Energy [EMSL-PNNL 51318 to J.C.D]. A portion of the research was performed using the Environmental Molecular Sciences Laboratory (EMSL), a DOE Scientific User Facility sponsored by the Office of Biological and Environmental Research and located at Pacific Northwest National Laboratory (PNNL).

## Author Contributions Statement

Conceived and designed the experiments: BW JJL JCD. Performed the experiments: BW XYZ ANK HDM LMM. Analyzed the data: BW XYZ ANK HDM LMM JJL JCD. Contributed reagents/materials/analysis tools: BW XYZ ANK HDM LMM. Wrote the paper: BW JCD. **Competing Interests Statement** The authors declare no conflicts of interests in the contents of this article.

## Figure Legends

**Supplementary Figure 1.**
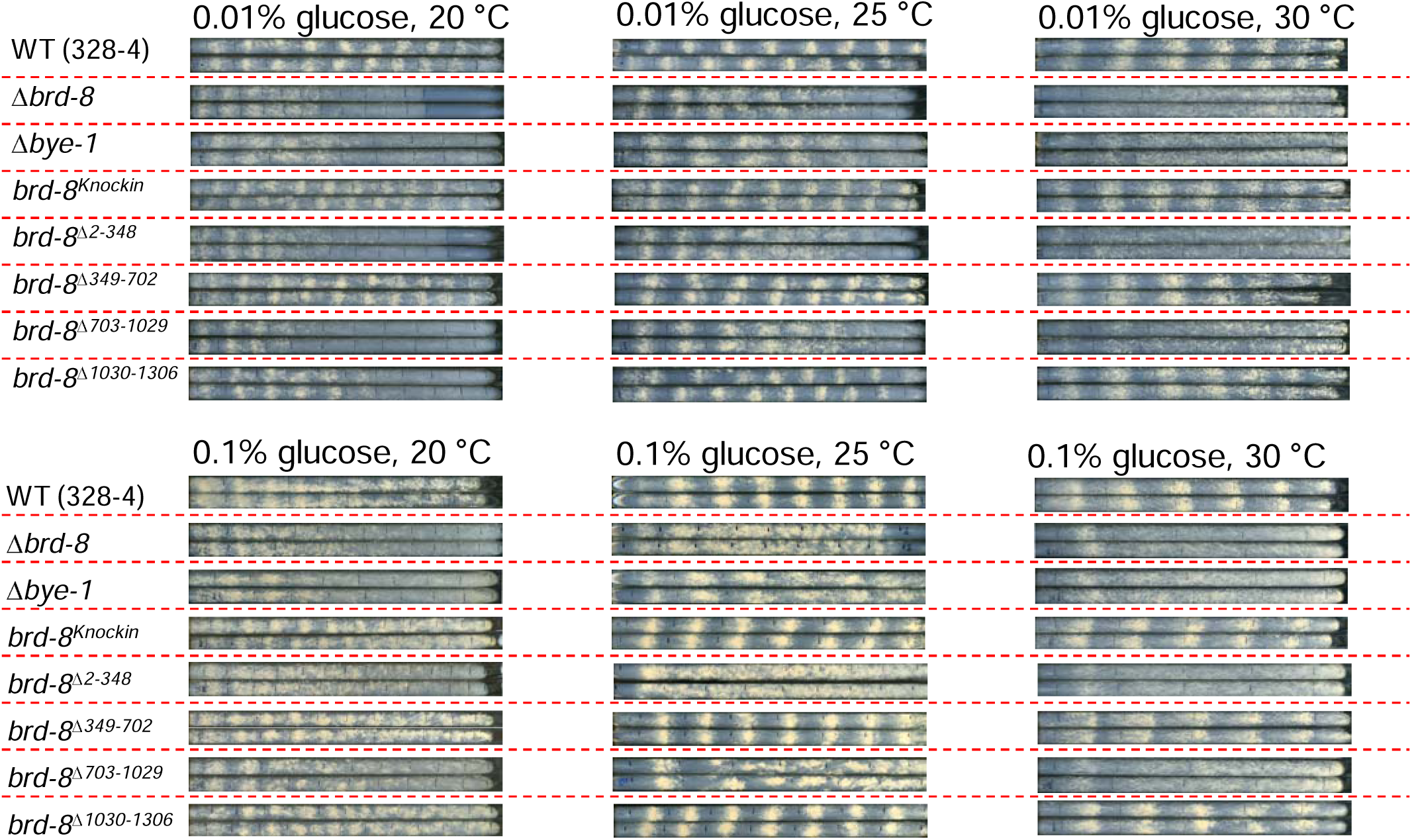
Race tube analyses of Δ*brd-8*, Δ*bye-1*, and *brd-8* mutants deleting indicated regions. The deletion or mutant strains were backcrossed to *ras-1^bd^* to facilitate visualization of circadian outputs. Race tube medium contains 0.1% or 0.01% glucose and cultures were incubated at 20, 25, or 30 °C as indicated. See Methods for details.

**Supplementary Figure 2.**
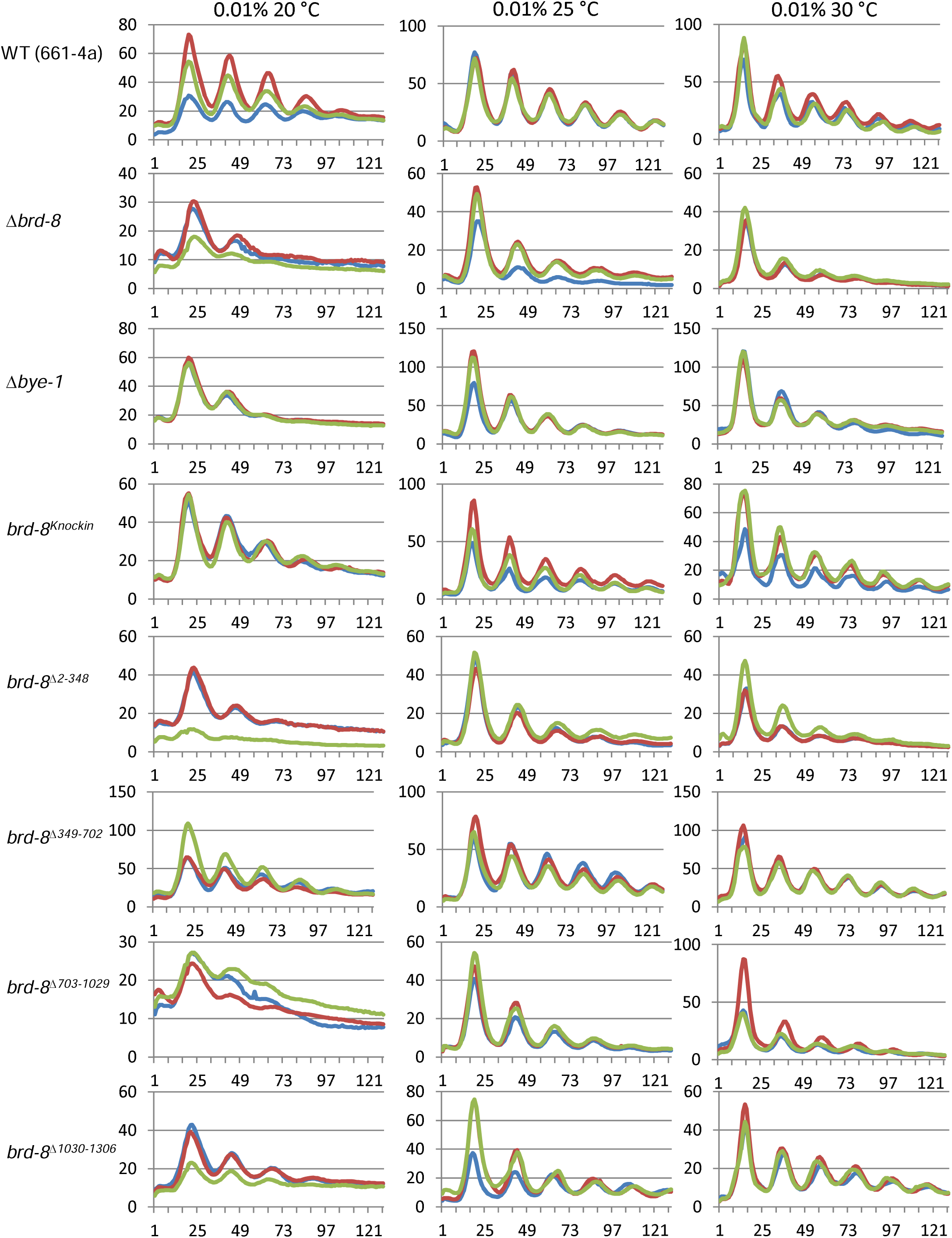
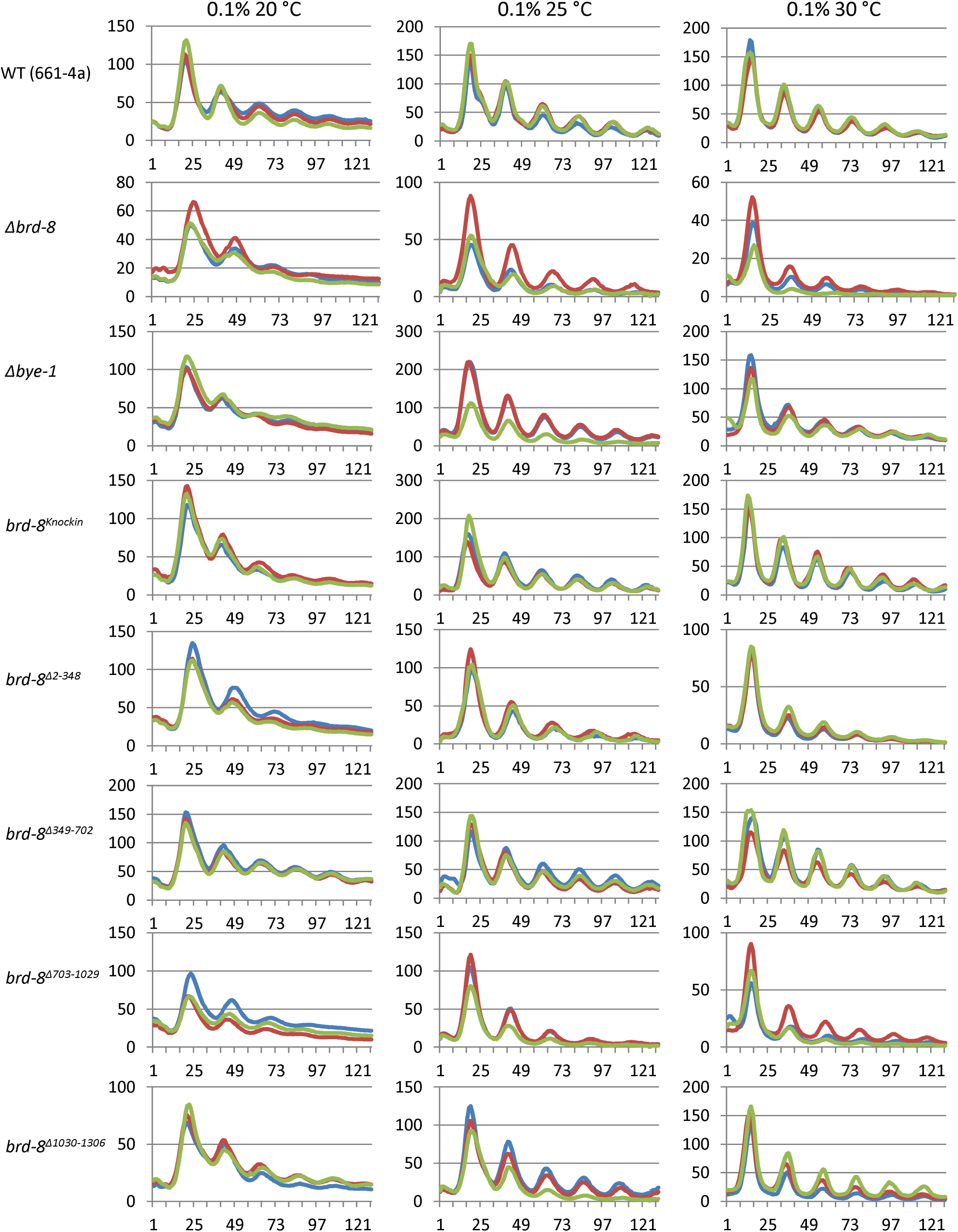
Luciferase analyses of Δ*brd-8*, Δ*bye-1*, and *brd-8* mutants. Strains were cultured in 96 well plates bearing race tube medium with 0.1% or 0.01% glucose and incubated at 20, 25, or 30 °C as indicated. Bioluminescent signals were recorded by a CCD-camera every hour, the data were obtained using ImageJ with a custom macro, and circadian period lengths were manually determined. Raw data from three replicates were plotted with the X-axis (time (hrs)) and Y-axis (arbitrary units) respectively.

**Supplementary Figure 3.**
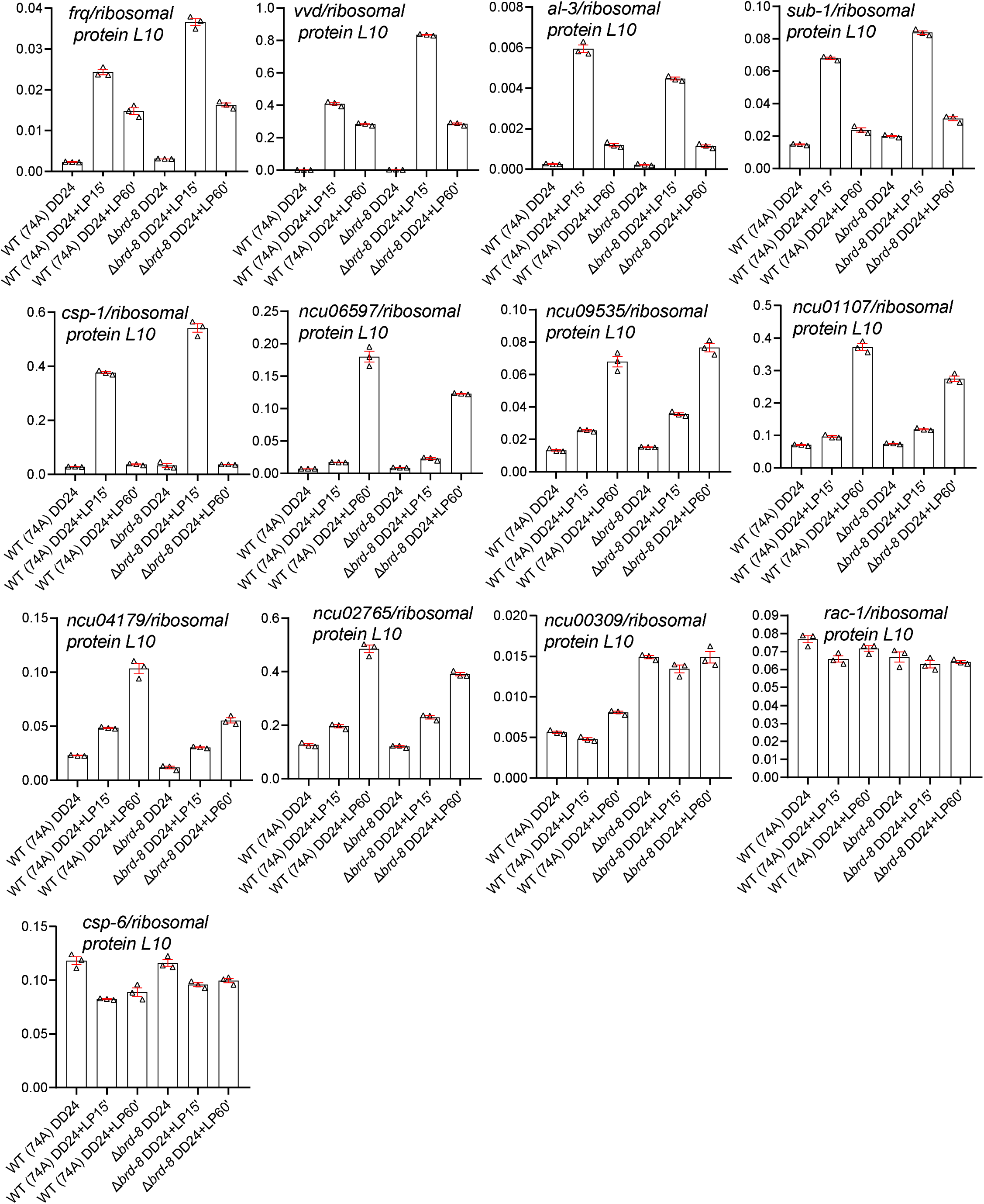
Expression of light responsive genes in WT and Δ*brd-8*. mRNA levels of indicated genes in WT and Δ*brd-8* were assayed by RT-qPCR with samples grown under dark for 24 hrs or treated with a 15– or 60-min light exposure after 24 hrs in the dark as indicated. Expression of these genes was normalized to that of ribosomal protein L10 (*ncu08964*). For RT-qPCR assays, three technical replicates were performed, and the bars represent average values from qPCR reactions for indicated genes normalized to those of *L10* (*ncu08964*), with error bars representing SEMs (n= 3). Source data were stored in the Source Data file.

**Supplementary Figure 4.**
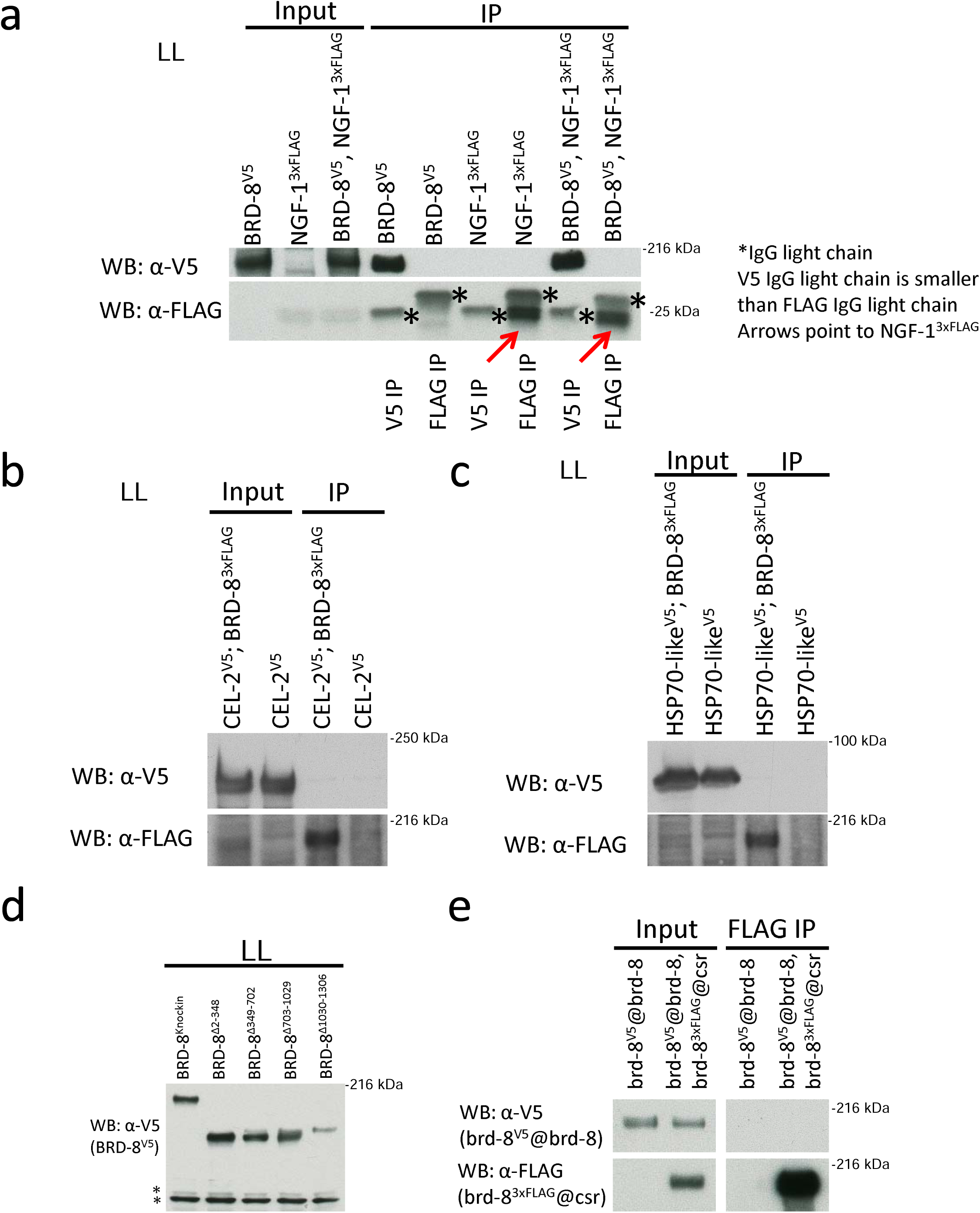
BRD-8^V5^ does not interact with NGF-1^3^ ^x^ ^FLAG^, CEL-2^V5^, HSP70-like^V5^, or itself (**a**) Immunoprecipitation was performed by FLAG or V5 antibody as indicated using BRD-8^V5^, NGF-1^3^ ^x^ ^FLAG^, and BRD-8^V5^/ NGF-1^3^ ^x^ ^FLAG^ followed by Western blotting against V5 or FLAG. Asterisks indicate IgG light chains of V5 or FLAG antibody and red arrows point to NGF-1^3^ ^x^ ^FLAG^. (**b**) CEL-2^V5^ (fatty acid synthase beta subunit dehydratase (NCU07307)); BRD-8^3^ ^x^ ^FLAG^ was immunoprecipitated with V5 and Western blotted by V5 and FLAG antibodies; CEL-2^V5^ serves as the negative control for the IP. (**c**) Heat shock protein 70-5 (HSP70-like, NCU08693) in BRD-8^3^ ^x^ ^FLAG^ was tagged with V5, immunoprecipitated with V5 antibody, and Western blotted with V5 and FLAG antibodies. HSP70-like^V5^ is the negative control for the IP. (**d**) Western blotting of *brd-8* deletion strains grown in the light at 25 °C verifying expression levels. (**e**) BRD-8 does not interact with itself. The native BRD-8 was tagged with V5 and a second copy of 3 x FLAG-tagged BRD-8 driven by its native promoter was knocked in the *csr* locus. BRD-8^3^ ^x^ ^FLAG^ was immunoprecipitated by FLAG antibody-conjugated resin and followed by Western blotting with V5 and FLAG antibodies respectively. Experiments in Supplementary Figures 4a–e were repeated twice (n = 3 in total), and similar results as the ones shown here were obtained. Source data except for the blots for Supplementary Figure 4a that were damaged in a lab fire were saved in the Source Data file. The marker position in Supplementary Figure 4a was estimated based on the molecular weight of the mouse immunoglobulin light chain (∼25 kDa), which was cross-reacted with the goat anti-mouse secondary antibody (IgG [Immunoglobulin G], HRP (horseradish peroxidase)-conjugated) in Western blotting.

**Supplementary Figure 5.**
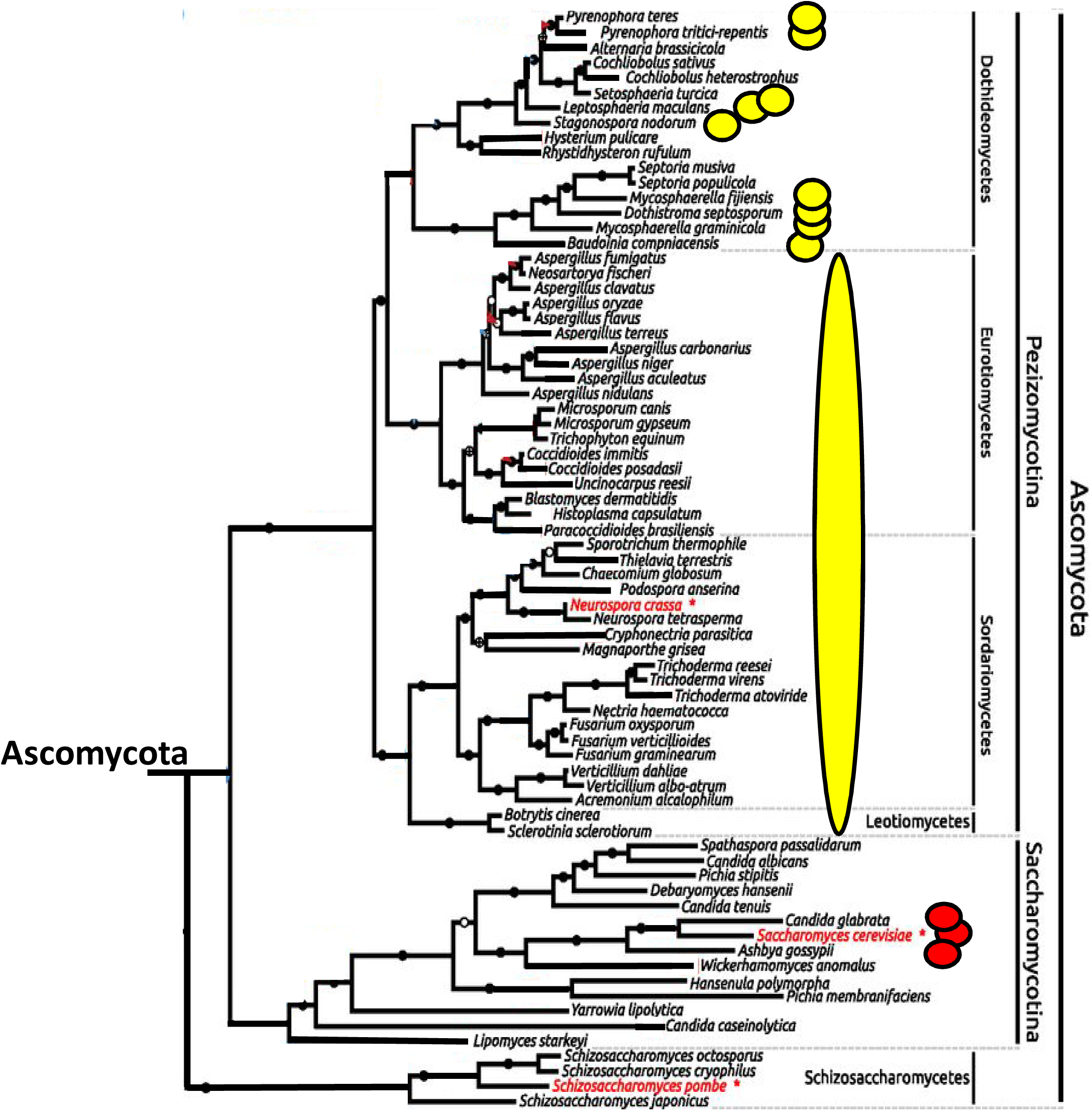
Phylogenetic conservation of *Neurospora* BRD-8 and Saccharomyces EAF5. Using a phylogenetic tree adapted from^96^, species having orthologs of BRD-8 are marked with yellow circles or ovals, and species with orthologs of EAF5 are marked with red circles. BRD-8 orthologs are universal within sequenced Leutiomycetes, Sordariomycetes, and Eurtiomycetes and common within Dothideomycetes, not all of which have sequenced proteomes available; EAF5 orthologs appear restricted to Saccharomyces and close allies. Orthology was determined by NCBI BLASTP searches as implemented on the SGD website [www.yeastgenome.org] using amino acid sequences for the two proteins.

**Supplementary Figure 6.**
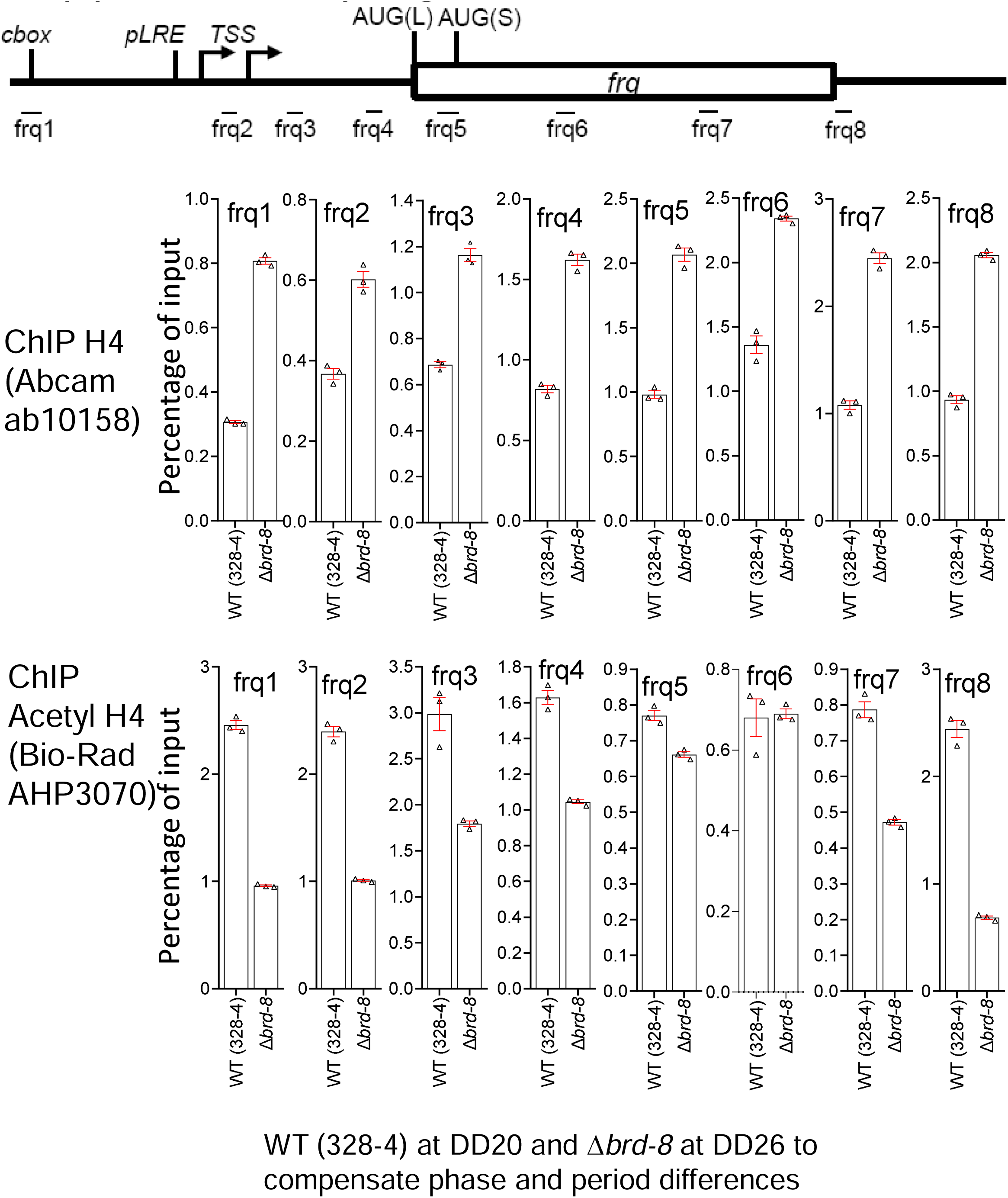
Validation of the ChIP-sequencing data of *frq* in Figure 4a by ChIP quantitative PCR assays. WT and Δ*brd-8* were harvested at DD20 and DD26 respectively after being crosslinked with formaldehyde for 15 min. ChIP experiments were done with antibodies against histone H4 or acetyl histone H4 (at K5, K8, K12, and K16) as indicated. The depiction of primer pairs used in the quantitative PCR reactions here is a copy of the one at the bottom of Figure 4a. See the Source Data file for source data.

**Supplementary Figure 7.**
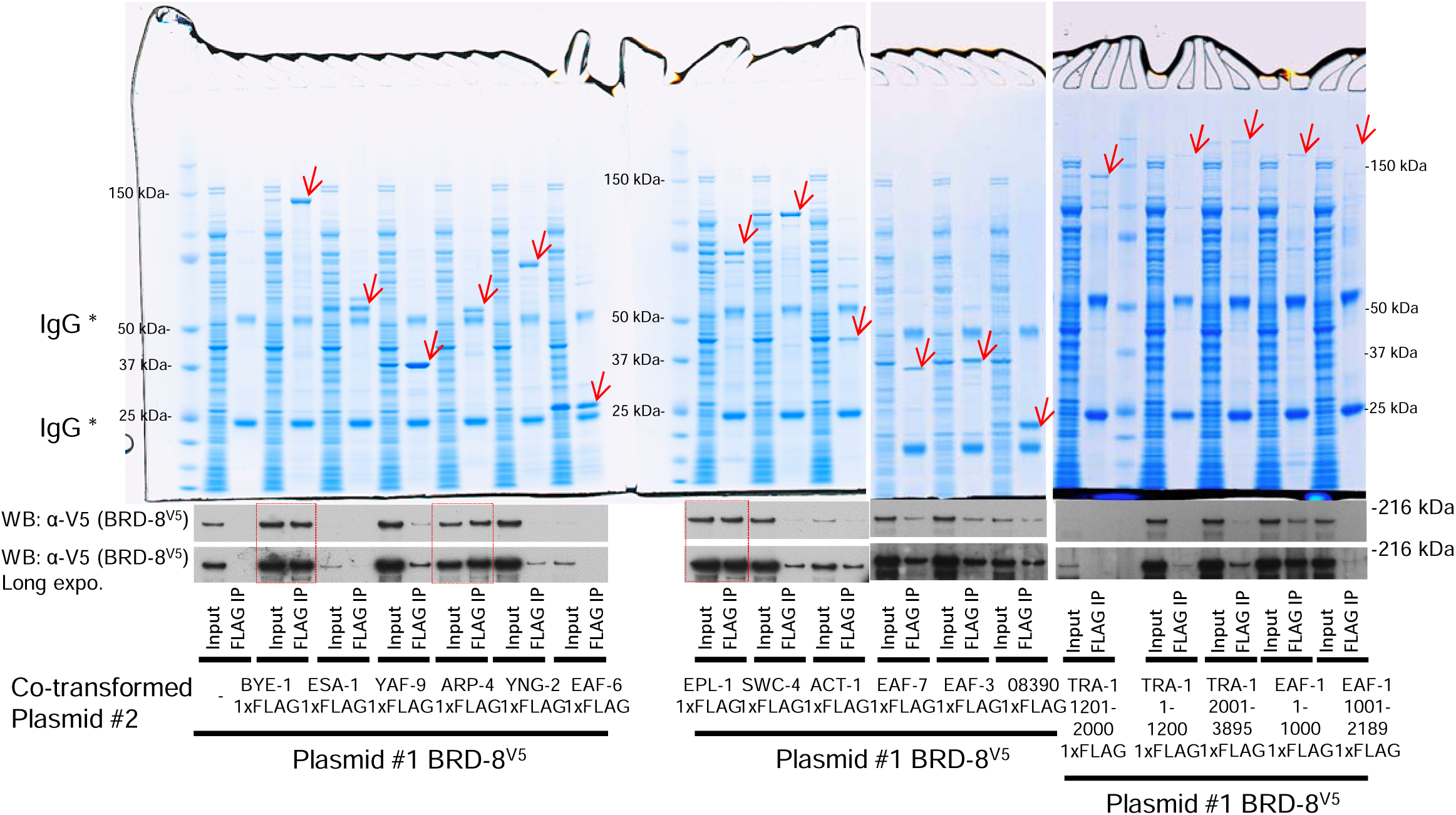
Interaction of BRD-8 and NuA4 subunits in bacteria. Plasmids expressing BRD-8^V5^ and BYE-1^1^ ^x^ ^FLAG^ or individual NuA4 subunits as indicated (tagged with 1 x FLAG respectively at their C-termini) were co-transformed to *Escherichia coli* (BL21 DE3), and their interactions were tested by IP with FLAG antibody. Gels were stained with Coomassie blue to show expression of BYE-1^1^ ^x^ ^FLAG^ and individual NuA4 subunits, and Western blotting was performed with V5 to display expression and interaction levels of BRD-8^V5^ with these proteins individually. Red arrows point to protein bands obtained from IPs with FLAG resin with the expected sizes; two exposures of V5 blots were shown to better visualize and compare BRD-8^V5^ levels in inputs and FLAG IPs. The assay in Supplementary Figure 7 was repeated three times independently with similar observations. Source data were included in the Source Data file.

**Supplementary Figure 8.**
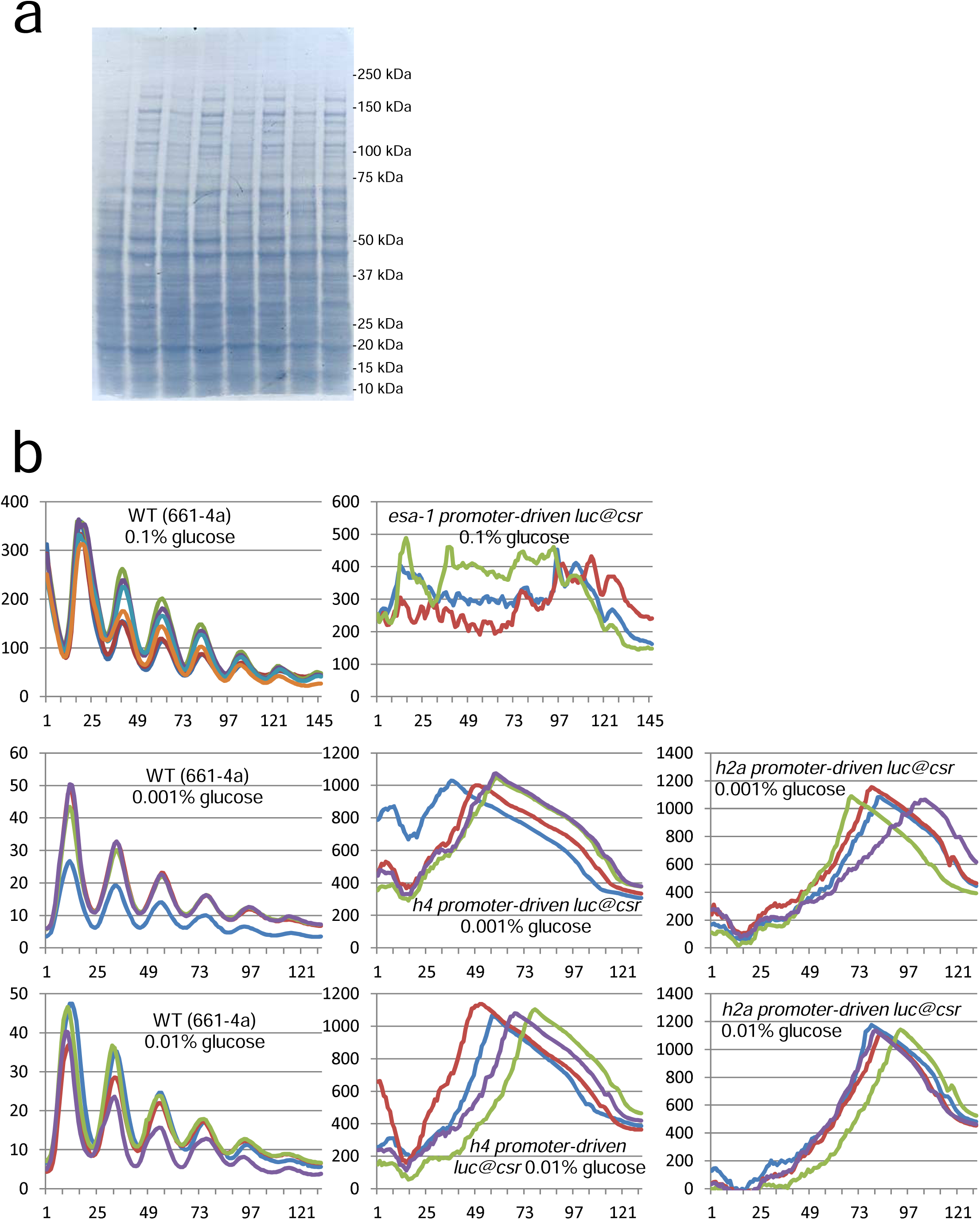
Promoter activity of *esa-1*, *histone h4*, or *histone h2a* in the dark. Blot staining for figure 6d (**a**) and luciferase analyses of the promoter of *esa-1*, *histone h4*, or *histone h2a* driving the *luciferase* gene at the *csr* locus (**b**). The assay was performed at 0.1, 0.01, or 0.001% of glucose in the culturing medium as indicated. The experiment in Supplementary Figure 8a was performed three times with similar results. Source data can be retrieved from the Source Data file.

**Supplementary Data 1** BRD-8^V5^ interactome identified by mass spectrometry. Cleared lysate from WT or BRD-8^V5^ was immunoprecipitated with V5 antibody-conjugated Dynabeads and the immunoprecipitated analyzed by mass spectrometry (See Methods for details).

**Supplementary Data 2** Densities of pan-histone H4 and acetyl histone H4 in WT at D20 hrs and Δ*brd-8* at D26 hrs. The six-hr lag between WT and Δ*brd-8* accounts for their period and phase differences between the two strains. The peak calling/comparison software SICER v1.1^97^ was used to assess differences in comparable peaks between WT and mutant samples. SICER assesses departure from a Poisson distribution to determine up-and down-regulated differences in peak size using separate one-sided tests; then an FDR correction is applied. Descriptions in details for the experiment can be found in Methods.

